# AXL governs axolotl cardiac regeneration and directs mammalian cardiomyocyte dedifferentiation

**DOI:** 10.64898/2025.12.21.695613

**Authors:** Elad Bassat, Jingkui Wang, Jorge Peña Peña, Inés Rivero-García, Agnieszka Piszczek, Francisco Falcon, Yuka Taniguchi-Sugiura, Jiaye Yang, Paula Fernández-Montes, Thomas Lendl, Lorena Domínguez, Katharina Lust, José Antonio Enríquez, Fátima Sánchez Cabo, Miguel Torres, Elly M. Tanaka

**Author notes:** Authors contributed equally (Elad Bassat, Jingkui Wang, Jorge Peña Peña). Corresponding authors (E.B.); (M.T.); (E.M.T.).

## Abstract

Cardiac injury outcomes vary widely across species, from complete regeneration to irreversible scarring. Using single-nucleus multiomics and spatial transcriptomics, we generated a spatially resolved atlas of axolotl heart regeneration following injury, identifying a distinct border-zone cardiomyocyte population with a pro-regenerative transcriptional program. Ligand-receptor analysis of the border zone niche identified enrichment of the receptor tyrosine kinase AXL in injury-responsive cardiomyocytes and its ligand Gas6 in endothelial cells. Functional perturbation demonstrated AXL requirement for border-zone cardiomyocyte activation and axolotl heart regeneration. In murine cardiomyocytes, AXL overexpression induced sarcomere disassembly, metabolic rewiring, and immature gene expression without triggering proliferation. These findings show that AXL signaling induces cardiomyocyte dedifferentiation and uncouples dedifferentiation from cell cycle re-entry, providing mechanistic insight into cellular plasticity during heart regeneration.

## Main

Cardiac diseases are among the leading cause of death worldwide (*1*). This is in part due to the inability of adult mammalian cardiac muscle cells (cardiomyocytes, CMs) to undergo effective dedifferentiation and subsequently proliferation and replenishment of the damaged myocardium (*2, 3*). In sharp contrast, regenerative species such as zebrafish and axolotls undergo epimorphic regeneration of their hearts throughout life (*4, 5*).

The extraordinary cardiac regenerative abilities of axolotls have been recognized for over half a century (*6*), well before similar capabilities were discovered in other species (*4*). Despite this early recognition, due to lack of available genetic and molecular tools, relatively little progress has been made toward understanding the underlying mechanisms of axolotl cardiac regeneration.

The processes underlying the regenerative response in zebrafish and neonatal mice have shed light onto the molecular mechanisms activated in CMs. These processes involve wide structural, metabolic and transcriptional changes, collectively known as dedifferentiation (*2*). This process in CMs is marked by several distinct hallmarks which take the cell from its differentiated, non-proliferative state to a less mature, expandable state. Structurally, the sarcomere is disassembled (*7–10*) and later reassembled with incorporation of reactivated immature sarcomere proteins (*11*) accompanied by cytoskeletal remodelling. Dedifferentiated CMs re-express fetal and developmental genes, as well as atrial and brain natriuretic peptides (Nppa, Nppb) (*12*). Metabolically, there is a shift from oxidative phosphorylation (Ox-Phos) to glycolysis, which is reminiscent of the fetal metabolic profile (*13, 14*).

The AP-1 transcription factor (TF) complex is an upstream regulator of these changes responsible for chromatin remodelling in genes related to regeneration (*8*). Inhibition of AP-1 activity in CMs impairs sarcomere disassembly, the invasion of the damaged area, and the proliferative and regenerative response (*8*). Other potent inducers of CM dedifferentiation include ErbB2 (*15*), Oncostatin M (*16*) and Yap1 (*17*), all of which induce sarcomere disassembly and re-expression of fetal genes. In all these cases, dedifferentiation is coupled to proliferation, which is frequently regarded as a core feature of the dedifferentiated state. Despite these advances, a critical knowledge gap remains regarding what governs the transition of homeostatic CMs to border zone CMs, what induces a robust and coordinated dedifferentiation program, and whether dedifferentiation alone is sufficient to drive a full proliferative response.

Here, leveraging the large size of the axolotl heart, which is similar in size to a mouse heart, we employed a combination of spatial transcriptomics and multi-omics (Fig. 1a) to characterize the axolotl cardiac cell populations and focused on changes occurring in the border zone CMs. We identified upregulation of Axl-Gas6 in the axolotl border zone following injury. In an axolotl cardiac cryoinjury model, dominant negative (dn) expression of AXL in CMs inhibited regeneration, altered border zone behaviour and moderately reduced CM proliferation. Adeno-Associated Virus (AAV) mediated AXL upregulation in mouse CMs promoted sarcomere disassembly and *in-vitro* a glycolytic shift in metabolism, as well as upregulation of immature-CM ion channel and border zone marker expression. Despite inducing these dedifferentiation events, AXL did not promote cell cycle activity in mammalian CMs. Overall, AXL is required for axolotl CM response to injury and is sufficient to induce *in-vitro* dedifferentiation of neonatal mouse CMs, thus having important implications for mammalian cardiac regeneration.

**Figure 1.**
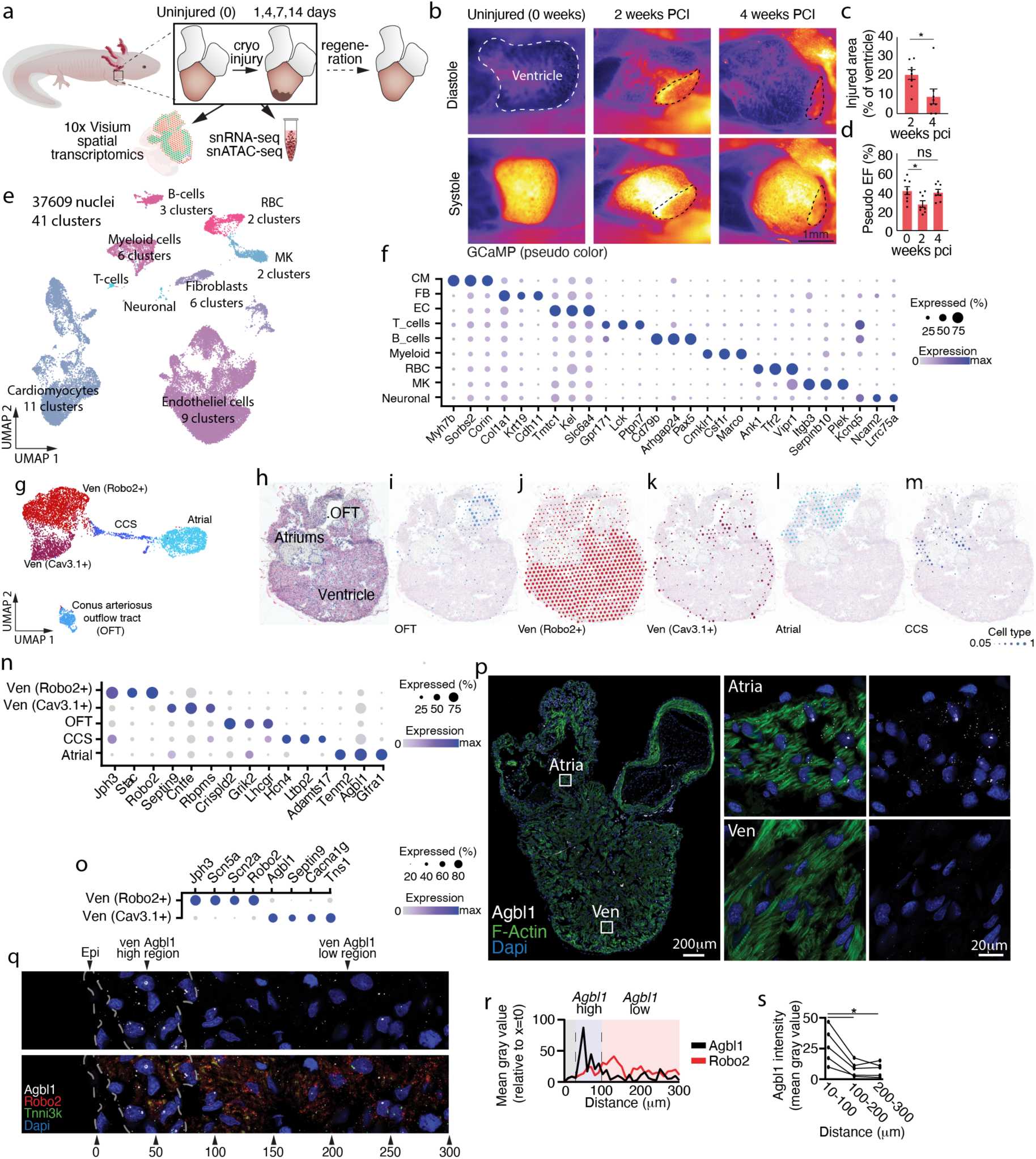
Cellular diversity of the Axolotl heart. **a**, Schematic representation of spatial transcriptomics and single nuclei multi-omics datasets collected in this paper. **b**, Representative images of Ca+ imaging in GCaMP6 transgenic axolotl hearts at peak systole and diastole, see also video s1-3. Dashed lines indicate the injury zone. **c**, Quantification of injured area size across time points. **d**, Pseudo-ejection fraction calculated from outline-based motion tracking of the heart in video recordings **e**, UMAP of snRNA-seq encompassing cells from 0,1,4,7 and 14 days after axolotl cardiac cryo injury. Each timepoint represents nuclei collected from 6 polled hearts. **f**, Dot plot depicting the top marker genes for each population. **g-m,** Sub clustering (**g**) and spatial localization of CM populations during homeostasis (**h-m**): OFT CMs (**i**), ventricular Robo2+ (**j**) and Cav3.1 (**k**) CMs, Atrial CMs (**l**), CCS (**m**) and in **n** the top marker genes for each sub population. **o**, dot plot depicting top marker genes when comparing Cav3.1+ and Robo2+ ventricular populations. **p**, HCR staining for Agbl1 focusing on atria and middle of ventricle. **q**, HCR staining for Agbl1 and Robo2 and Tnni3k in the ventricle and quantification of agbl1 in **r,s**. In **<u>s</u>**, representative average expression based in **t**. In **s**, each line represents quantification of individual heart. \**P* < 0.05, \*\***P** < 0.01, \*\*\***P** < 0.001 (statistical test: one-way Anova with Geisser-greenhouse correction and Tukey’s post hoc).

### Axolotl cardiac injury size peaks after two weeks and induces cardiomyocyte cell cycle activity

We performed a time course analysis after cryoinjury in hearts of 10 cm axolotls. We observed significant injury areas peaking at 2 weeks following cryocauterization, with progressive reduction in the injury area during the following weeks (Fig. S1a,b). In accordance with previous salamander and zebrafish studies (*5, 18, 19*), some injured area was evident even after 8 weeks, and likely would require 4 months for complete regeneration (*20*). The same trend of reduction of injury size between 2 and 4 weeks was also seen by using Ca+ reporter animals in which GCaMP6 becomes fluorescent upon Ca+ binding (Fig. 1b-d, vid. 1a-c). As seen in other regenerative species (*7, 10*), axolotl cardiac injury induces a marked increase in CM cell cycle activity (*5*). Using both EdU (Fig. S1c,d) and a fluorescent ubiquitination-based cell cycle indicator (Fucci) (*21*) (Fig. S1e,f), we found significant cell cycle activity peaking at day 7 after injury and persisting after 3 weeks.

### Cellular diversity of the axolotl heart

In order to identify and characterize the cells of the axolotl heart we utilized an integrative transcriptomic strategy using single-nucleus RNA sequencing (snRNA-seq) and single-nucleus assay for transposase-accessible chromatin sequencing (snATAC-seq) from the same cell (multiomics). To annotate the cell types in the homeostatic and regenerating heart, we performed multiomic analysis of uninjured and cryoinjured at the following stages: non-injured (d0), inflammatory phase (d1), proliferative phases (d4/7) and beginning of maturation stage (d14), based on mammalian dynamics (*22*) (Fig. 1e, Fig. S2).

We ultimately recovered 37,690 nuclei and were able to identify 9 major clusters which could be divided based on canonical markers shared with other species such as Tnni3K for cardiomyocytes (CM), Pecam/*Cd31* for endothelial cells (EC), *Col1a2* for fibroblasts (FB), *Itgam* for Myeloid cells, *Pax5* for B-cells and *Mpl/Itgb3* for megakaryocytes/thrombocytes (MK). From that we could define the top 3 markers for each major cluster: CM (*Myh7b/Sorbs2/Corin*+), EC (*Tmtc1/Kel/Slc6a4*+), FB (*Col1a1/Krt19/Cdh11*+), myeloid cells (*Cmklr1/Csf1r/Marco*+), B-cells (*Cd79b/Arhgap24/Pax5*+), red blood cells (RBC, *Ank1/Tfr2/Vipr1*+), MK (*Itgb3/Serpinb10/Plek*+), T-cells (*Gpr171/Lck/Ptpn7*+) and neuronal (*Kcnq5/Ncam2/Lrrc75a*+) (Fig. 1e,f Fig. S3a). Although we could account for most major cell types of the heart, we could not definitively identify smooth muscle cells, as key markers were expressed in different populations, such as *Myh11* exclusively in the megakaryocytes/thrombocytes cluster (data not shown). Given the lack of developed coronary vasculature in amphibian hearts (*23*), it is possible that the heart lacks such cells. Further clustering of cells identified a total of 41 unique sub-clusters for which we could define unique characterizing markers, including at least one specific proliferative population for each major cluster (Fig. S3b,c). Out of the 41 identified clusters, 11 were injury-specific populations (Fig. S3d).

To further gain spatial information of those annotated cell types, we next used a spatial transcriptomic approach (10x genomics Visium) on similar samples and time points. After performing manual segmentation of border and remote zones on replicates from day 0,4 and 7 after injury, the Visium data showed segregation based on the localization while showing almost no segregation based on replicates suggesting the high reproducibility of our data (Fig. S3e,f). Using robust cell type decomposition (RCTD) (*24*) we determined the localization of the individual cell types by “deconvolving” the Visium 55um pixel, essentially mapping the single nuclei data to the spatial data.

Focusing on homeostatic (d0) hearts, the fibroblast population showed the lowest diversity, both in regards of number of cells as well as number of unique clusters (Fig. S4a,b). From the 3 annotated clusters, *Pkd1*+ population was the largest and expressed Complement Factors, *Efemp1*, various keratins and *Bnc1* all commonly expressed by epicardial cells (Fig. S4b,c) (*25, 26*). Mapping *Pkd1*+ population onto the Visium slide showed that indeed they are mostly found in the periphery of the heart, suggesting an epicardial localization (Fig. S4d). This was further validated using *in-situ* hybridization chain reaction (HCR) targeting *Pkd1*+ cells (Fig. S4e). The two other FB clusters were enriched for ECM proteins and marked by *Tnxb/Lama2/Ngf*+ and *Vwa2/Cspg5/Acan*+ (Fig. S4b). Mapping the *Tnxb*+ and *Vwa2*+ populations onto the Visium slide showed an interesting localization pattern, as *Vwa2+* population was found exclusively in the area between the atria and ventricle, adjacent to the location of the atrio-ventricular canal, as well as in the conus arteriosus outflow tract (OFT) (*27*) (Fig. S4f). The *Tnxb*+ population was surrounding the *Vwa2*+ population in both locations as well as to a lesser extent found in the periphery of the heart (Fig. S4g). The localization of the *Vwa2*+ population in the base of the ventricle is found within a low muscle density region which shows some ECM rich characteristics, correlating with the high expression of ECM genes in this population, making them likely to be related to annulus fibrosus and valve fibroblasts.

Endothelial cells were the most abundant cell type in the heart, in accordance with their relative abundance in other species (*28*), despite the lack of ventricular coronary plexus in axolotls (*23*). In steady state, ECs distribute in 5 different clusters with unique transcriptomic signatures (Fig. S4h,i). The largest population, which we termed “main”, showed high levels of *Lama2/Kcnmb2/Radil*+, however these marker genes were not unique but rather showed varying expression levels of genes shared with the NOS3+ cluster, suggesting that these populations are related to each other and possibly represent different functional states of the same cell type. Spatial localization of the EC_main and Nos3 populations showed specificity to the ventricle with minimal expression in other chambers (Fig. S4j,k). *Pth1r*+ population showed high expression of *Prox1*, usually indicative of endothelial lymphatic cells, although also found in valvular endothelial cells (*29*); however, as our Visium sections did not clearly include valves, we could not assign this population. The *Lhx6*+ population showed high expression of *Flt4* and *Lyve1* (Fig. S4l), both markers of lymphatic endothelium(*30*), however it is currently unknown whether lymphatic vasculature, even in an immature form, exists in the axolotl heart. In the ventricle the *Lhx6*+ and *Pth1r*+ endothelial populations are scarcely spread throughout and do not restrict to specific localizations. In general, when comparing the markers of different EC clusters those described in other model organisms, no clear cell type homology was discernible.

CM diversity in the absence of injury was captured in 5 distinct clusters (Fig. 1g), all of which showed high levels of sarcomere protein-coding genes (Fig. 1f, Fig. S3a). In order to sub-cluster the CM populations, we looked at the location of the cells in the spatially resolved dataset and saw that each cell type has a unique localization, which allowed us to annotate them based on geographical localization (Fig. 1h-m). One population mapped onto conus arteriosus (outflow tract: OFT) CMs (Fig. 1i), two were ventricular populations (*Cav3.1*+/ and *Robo2*+, as marked by one of their top differentially expressed gene, DEG) (Fig. 1j,k), one atrial population (Fig. 1l), and a population of the cardiac conduction system (CCS) (Fig. 1m). The CCS population was mainly identified by expression of a known marker, *Hcn4*, and its distinct spatial distribution compatible with the possible locations of sinoatrial and atrioventricular nodes (Fig. 1m) (*31*). Using these annotations, we could identify top DEGs for each population: OFT (*Crispld2/Grik2/Lhcgr*+), Atrial (*Adamts17/Agbl1/Gfra1*+), CCS (*Hcn4/Ltbp2/Tenm2*+), ventricular (v) CM_*Robo2*+ (*Jph3/Stac/Robo2*) and vCM*_Cav3.1*+ (*Septin9/Cntfr/Rbpms*) (Fig. 1n). To better distinguish the two ventricular populations, we analysed the DEGs specifically between these two populations. We could identify that the CM_*ROBO2*+ population expresses two Sodium Voltage-Gated Channels: *Scn5a* and *Scn2a*, which encode for Nav1.5 and Nav1.2, respectively. Both channels are expressed at low levels in the CM_*Cav3.1*+ population (Fig. 1o). The Cav3.1 population showed high levels of *Cacna1g*, a gene encoding for a subunit of a voltage-dependent Calcium channel. *Agbl1* was identified as differentially expressed between the CM populations, being predominately expressed in the atria (Fig. 1n), while also differently expressed between ventricular populations (Fig. 1o). HCR staining against *Agbl1* corroborated the gene expression data, as it was highly expressed in the atria with lower expression in ventricle (Fig. 1p). Focusing on the ventricle, *Agbl1* expression was identified to also distinguish between the two ventricular populations, with higher levels of expression in the *Cav3.1*+ population (Fig. 1o). HCR staining for *Agbl1* showed expression mostly in the periphery of the heart proximally to the epicardium (Fig, 1q-s), in accordance with the predicted population localization (Fig. 1j,k). Collectively, we defined 41 clusters, out of which 30 are found also in the absence of injury, from this we generated a list of DEG identifying markers for each of these clusters.

### Spatial analysis of interstitial heart cells after injury

Having obtained the spatial localization of the cardiac cell populations, we next wanted to analyze the composition of cells after injury. For that, we focused on populations that are both sufficiently abundant and localize to the ventricle, the region which was injured.

Early after injury we observed the emergence of a transient epicardial cell population that still expresses epicardial markers but differs from the uninjured epicardium by ±1000 genes from which the most significantly upregulated are *Has1/Tfpi2/Cxcl14* (Fig. S5a,b). By day 4 after injury the *Tfpi2*+ cluster is largely missing and the *Pkd1*+ population returns (Fig. S5c). The upregulation of *Has1* suggests increased hyaluronic acid (HA) synthesis after injury and, although this gene has not been previously shown to be upregulated after cardiac injury in the epicardium, the role of HA in regeneration has been extensively documented (*32*). Of note, we could not identify any of the canonical “activation” or EMT-related epicardial markers upregulated in this population. On the contrary, the embryonic epicardial gene *Tbx18* which is usually upregulated after injury was downregulated in the injury specific population suggesting that the axolotl epicardial response to injury differs from that in other species(*33*). The major changes occurring in EC after injury were the reduction of the *Pth1r*+ population and replacement with the *Iars1*+ population 1 day after injury and then at day 4 the appearance of a persistent *Lox*+ population which remains throughout our time course. An additional proliferating EC population persisted in all time points (Fig. S6a-c). The *Iars1*+ population showed localization mostly in the infarct region whereas the *Lox*+ population was rather found to be more dispersed in the border zone and surrounding viable tissue as well as the injury zone (Fig. S6d). All the injury-specific populations expressed high levels of *Serpine1* (Fig. S6e), which was previously shown to be enriched in the wounded endocardium and play an important role in both EC and CM proliferation(*34*). Trajectory analysis using Partition-based graph abstraction showed a distinct network for the homeostatic cell types and for the injury specific/proliferating cell types, which is connected by the main EC population (Fig. S6f). Further analysis of the injury-specific populations showed that the main EC population contributes to the injury specific *Iars1*+ population, which later contributes to both the injury-specific *Lox*+ and the injury-specific proliferating population (Fig. S6g).

The cardiac immune cell populations were mostly comprised of myeloid cells (Fig. S7a,b). To annotate the myeloid sub-clusters, we performed sub-setting based on *Cd45/Itgam*+ expression. Macrophages were identified based on the expression of *Csf1r/Cd68*+ which are conserved also in other species. Due to lack of Ly6g expression outside of mammals, we relied on common markers from zebrafish, mainly myeloperoxidase (*Mpo*), *Mmp9* and *Csf3r* to annotate neutrophils (*35, 36*) (Fig. S7c,d). Using these markers, we identified a small neutrophil population marked by top DEG *Csf3r/Arg1/Il1r1*+ which also expresses *Mpo* and *Mmp9* (Fig. S7d). Using this methodology, we identified 6 different myeloid clusters; two of which are found during steady state at low numbers: resident monocytes/macrophage_1 (Mo/Macs) (*Cd163l1/Axl/Adgrl3*+) and resident Mo/Macs_*dysf*+ (*Hvcn1/Cfp/Dysf*+). Other recruited Mo/Macs populations are marked by *Faxdc2/Plbd1/Parvg*+, *Snx22/Itgad/Lgals9*+ (Fig. S7c).

During steady state, low numbers of resident monocytes/macrophage (Mo/Macs) were found dispersed in the heart. 1 day after injury, resident *Dysf*+ and *Snx22*+ recruited Mo/Macs began to accumulate and were seen dispersed throughout the ventricle (Fig. S7e,f). In addition, the neutrophil population reached their peak numbers and were largely localized to the injured and border zone (Fig. S7e,f). At 4 days the peak of Mo/Macs recruitment was seen with continued accumulation of *Snx22*+ and appearance of *Faxdc2*+ population, as well as dwindling yet persistent numbers of the *Dysf*+ resident population (Fig. S7e), all of which were localized in the vicinity of the injury and border zone. After 7 days the *Faxdc2*+ was the prevalent myeloid population of the heart and the *Snx22*+ largely disappears. After 14 days most neutrophils were already cleared from the heart and, except for residual *Faxdc2*+ cells, most recruited Mo/Macs were removed from the heart and only the resident populations persisted although with higher immune cell numbers compared to non-injured time (Fig. S7e). Comparing these Myeloid populations to zebrafish populations did not show any particular trend, as markers identified by specific ZF populations (*37*) were largely absent in axolotl cells or were shown to be dispersed on multiple populations. Axolotl resident monocytes/macrophage population showed high expression of *Timd4* (Fig. S7g), a known marker of resident macrophage population in mice, however they also showed expression of *Ccr5* (the closest *Ccr2* homeolog in axolotls) which is not commonly expressed in the same cells as *Timd4* (*38*).

Taken together cardiac interstitial cells show dynamic changes after injury. Epicardial cells and endothelial cells responded quickly and already showed distinct populations 1 day after injury, however they diverged in the extent to which the responsive population persisted. The myeloid populations followed a similar trajectory to those of other species (*22*) in which an early response of neutrophils was observed which was gradually replaced by macrophage populations. Of note, a resident macrophage population, existing also during steady state, persisted in the heart throughout the injury time course and showed transcriptomic proliferative signature similarly to self-maintained macrophage population in other species (*38*). However, unlike mammalian resident macrophages, which were greatly reduced in numbers after injury before replenishing (*38*), the axolotl resident macrophage population was maintained throughout the time course.

### Dynamic changes in cardiomyocytes following cardiac injury

CMs displayed region-specific responses to injury, with ventricular populations showing the most dynamic changes. OFT CMs remained unaffected, while atrial CMs showed only a transient response marked by a short-lived *Pdcd6ip*⁺ population (Fig. S8a,b). In the ventricle, during homeostasis two major clusters were identified: Robo2⁺ and Cacna1g (Cav3.1+) along with two proliferative clusters (Prol_1 and Prol_3) enriched in cell cycle genes (upregulating Aspm/Cenpf/Anln⁺ and Enf1/Cenpe/Diaph3⁺ respectively, Fig. S8b). Following injury, Cav3.1+ cells decreased while a new injury specific (CM I.S.) population appeared. By days 4-7 CM I.S. declined while CM Prol I.S. increased until day 14 (Fig. 2a).

**Figure 2.**
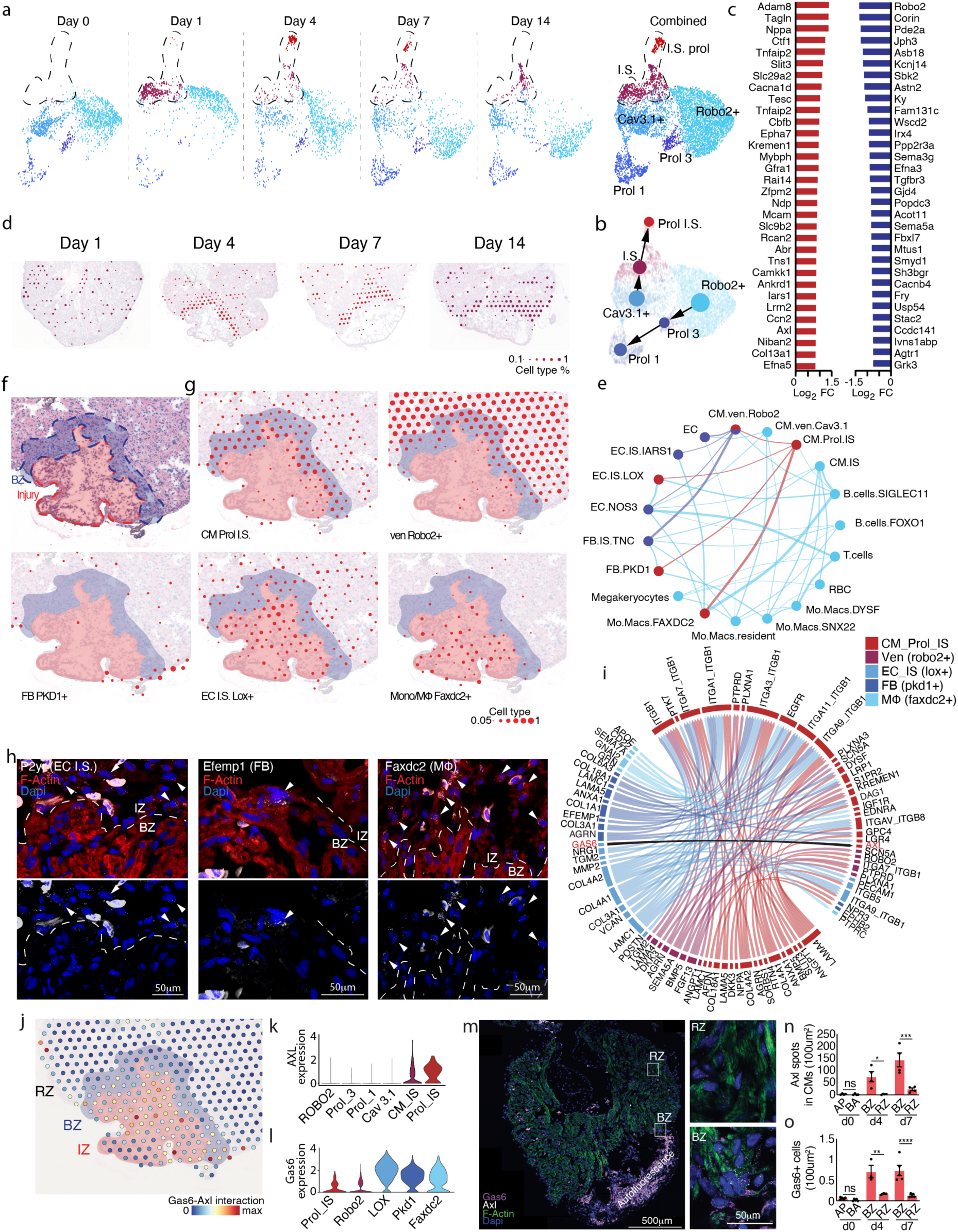
Cellular diversity of cardiomyocytes after injury. **a**, UMAP depicting the temporal changes of ventricular CM populations. Dashed line focuses on the injury specific populations. **b**, Partition based graph abstraction (PAGA) trajectory analysis of the CM populations predicted two distinct trajectories one of which is injury specific. **c,** Top up and downregulated genes for the I.S. prol population. **d**, Spatial mapping of the I.S. (day 1 and 14) and I.S. Prol (day 4 and 7) populations on the H&E stained ventricle section. **e**, Spatial proximity network of all cell subclusters present within the ventricle at day 4 after injury with MISTy neighborhood analysis. Red nodes and lines represent the cell clusters with higher score of colocalizing with the CM.Prol.IS (border zone). Blue nodes and lines represent the cell clusters with higher score of colocalizing with the homeostatic Robo2+ population. Half-red and half-blue nodes are shared between the BZ and homeostatic cells. Thicker lines represent higher colocalization scores. **f**,**g** Enriched localization with BZ cell type overlay on H&E stained visium slide showing all identified types are found within the BZ (mask in blue, as inferred by CM.Prol.IS localization profile). Dots represent each visium “pixel”. Size represents the proportion of the specific assayed cell type within the pixel. **h**, HCR staining against key markers of each identified enriched cell population showing localization within the BZ and adjacent regions. Arrowheads show HCR positive cells. **i**, Day 4, top 100 LIANA ligand-receptor interaction map of the cell types found proximal to BZ CM (from panel **a**). Top half represents the receptors; bottom half represents the expressed ligands. Arrow colors represent the ligand sending cell. Black arrow shows Gas6-Axl interaction. **j**, Gas6-Axl Ligand-receptor interaction overlaid on a H&E stained slide, 4 days after injury. IZ-injury zone, BZ-border zone, RZ-remote zone. **k,l,** Violin plot showing Axl expression in the ventricular CM populations (**k**) and Gas6 expression in the cells encompassing the BZ niche (**l**). **m**, Representative HCR staining of day 4 heart after injury probing for Gas6 and Axl with counterstain of Phalloidin and Dapi. Squares depicting magnified regions on the right of RZ (remote zone) and BZ (border zone). **n**, HCR quantification of the number of HCR-Axl spots per CM within the border zone. AP, apical side of the heart. BA, basal side of the heart. **o**, HCR quantification of the number of Gas6 positive cells adjacent to the BZ or RZ. \**P* < 0.05, \*\***P** < 0.01, \*\*\***P** < 0.001 ****P<0.0001 (statistical test: two-way ANOVA with Bonferroni post hoc).

Transcriptomic and trajectory analyses uncovered two divergent pathways, distinguishing homeostatic maintenance from injury-induced responses. Cell cycle scoring based on known cell cycle regulators expression confirmed that Prol 1 and Prol 3 were highly proliferative, clustering according to cell cycle phase (Prol 3-G2M+, Prol 1-S+, Fig. S8c). The injury specific populations showed a mix of both S and G2M phases starting at 4 days persisting till 14 days after injury (Fig. S8c). Using Partition-based graph abstraction (PAGA) (*39*) with RNA velocity showed two distinct trajectories: *Robo2*+ → Prol 3 (S) → Prol 1 (G2M) which we term the “maintenance” trajectory (present in homeostasis) and a *Cav3.1*+ → I.S. → Prol I.S. “injury” trajectory. Of note, no contribution from *Robo2*+ to Cav3.1+ (or vice versa) could be seen. The unidirectional Prol 3 → Prol 1 (i.e. S → G2M) transition confirmed biological plausibility, though the post-cycle fate of cells remains unclear.

CM Prol I.S. showed >6000 DEGs compared to homeostatic populations, including conserved “border zone” markers, *Nppa* and *Ankrd1* (*40*) (Fig. 2c), as well as *Ctf1* and *Ccn2* (*Ctgf*), both linked to CM protection, proliferation and tissue repair (*41, 42*). Notably, CM Prol I.S. population also expresses the receptor tyrosine kinase AXL, previously implicated in proliferation, migration, survival of cells (*43*) and recently also in oral regenerative wound repair (*44*), but never in CMs or in the context of cardiac regeneration. Ingenuity pathway analysis (IPA) of predicted upstream regulators showed known facilitators of cardiac regeneration such as: MYC (*45, 46*), NRG1 (*47*) and ERBB2 (*15*) (Fig. S8d) to be upregulated in Prol I.S. population. As the injury specific populations are proliferative and show hallmarks of BZ CMs, we wanted to check whether they are indeed localized to the border zone. Mapping the CM I.S. and CM Prol I.S. population, based on their abundance in the trajectory (i.e. CM I.S. expressed on days 1 and 14 while CM Prol I.S. on days 4 and 7) on the spatially resolved dataset showed that it maps in the 50-200um (1-3 Visium spots) surrounding the injury, suggesting that they are indeed border zone CMs (Fig. 2d). HCR against known BZ markers *Nppa* and *Ankrd1* confirmed the localization to the border zone (Fig. S8f,g).

Next, we wanted to identify which transcription factors might control the transition from homeostatic to injury specific populations. Using our snATAC-seq, we applied motif enrichment analysis with chromVar and identified strong enrichment for AP-1 family factors (19 of the top 20 motifs; Fig. S8e) in Prol I.S., consistent with previous reports in border zone CMs (*8, 40*).

Together, these data suggest that the ventricle possesses two distinct homeostatic populations, one sustaining a maintenance\growth proliferative capacity and a second driving an injury specific response. These injury-specific cells are localized to the BZ, and have a unique transcriptomic signature, upregulating canonical BZ genes such as *Nppa* and *Ankrd1* as well as novel genes such as *Axl*.

### Spatial organization and signaling at the injury border zone

After obtaining spatially resolved datasets and annotating the different cell types based on temporal and spatial context, we next examined the degree of direct interactions between cell types based on their colocalization within a Visium pixel. For that, we employed Multiview Intercellular SpaTial modeling framework (MISTy) neighborhood analysis (*48*) and evaluated regions of two sizes: Within the same Visium pixel (55um) and within adjacent Visium pixels (100um between pixel centers). These 2 different region sizes also stratified the potential of cell types for direct interaction, given closer cells have a higher probability of interacting. Focusing on the CM populations, in the uninjured heart (day 0), *Robo2*⁺ CMs showed no strong spatial association with other cell types, whereas *Cav3.1*⁺ CMs, localized to the heart periphery, interacted with epicardial (*Pkd1*⁺), endothelial (EC), and *Robo2*⁺ CM populations (Fig. S9a). After injury, both injury-responsive and homeostatic *Robo2* CMs display new interaction networks. By day 1 post-injury, both populations colocalized with *Nos3*⁺ ECs, injury-responsive *Iars1*⁺ ECs, *Tfpi2*⁺ fibroblasts (FBs), and *Snx22*⁺ macrophages (Fig. S9b). As the injury response progressed, we observed an overall increase in interaction complexity, with a peak at days 4 and 7, followed by a decline by day 14, coinciding with the gradual return of most cell types to a homeostatic transcriptomic state. Starting from day 4, a clear pattern emerged: border zone CMs consistently interacted with at least one population from each of three key lineages: immune cells, ECs, and FBs/epicardial cells, though the specific subpopulations varied across time. At day 4, homeostatic CMs remained associated with *Nos3*⁺ and *Iars1*⁺ ECs and began interacting with *Tnc*⁺ FBs, while CM.prol.IS cells localized with *Lox*⁺ ECs, *Faxdc2*⁺ macrophages, and epicardial cells at the periphery of the border zone (Fig. 2e–h). These interactions were confirmed by HCR staining for representative differentially expressed genes (Fig. 2h). By day 7, both CM populations retained interactions with ECs but diverged in immune and stromal contacts: homeostatic CMs localized with resident *Dysf*⁺ macrophages, while CM.prol.IS cells associated with *Tnc*⁺ FBs and *Snx22*⁺ macrophages (Fig. S9c). At day 14, this tri-lineage interaction pattern persisted, albeit with reduced overall complexity, with *Robo2*⁺ CMs weakly colocalizing with resident macrophages, and CM.IS cells associating with T cells and *Dysf*⁺ macrophages, while both retained EC interactions (Fig. S9d). Although global transcriptomic profiles at days 4 and 7 appeared similar between biological replicates (Fig. S3e,f), the specific cellular compositions of the BZ niches varied. This variation suggests a highly dynamic and plastic regenerative environment, where multiple immune, stromal, and endothelial subtypes may engage with BZ CMs in different replicates. For subsequent analysis, we focused on one representative network, while acknowledging that other interactions likely play equally relevant biological roles.

Focusing on the niche found surrounding the border zone, we next wanted to understand the communication between the adjacent cell types by utilizing LIANA (*49*), that models intercellular interactions by linking ligands and receptors. In the generated L-R interaction list, we identified many previously known interactions involved in cardiac response to injury, such as AGRN-DAG1 (*50*) (Fig. 2i), NRG1-ERBB2:ERBB3 (*15*) (data not shown, found in top 200 interactions when only top 100 are shown), as well as novel interactions which were never discussed in this context, such as AXL-GAS6 (Fig.2i). Mapping the *Gas6-Axl* pair on our spatially resolved dataset showed that they are particularly enriched in the border zone compared to the remote area (Fig. 2j). In addition to the border zone, the mapping also showed high level of interaction in the injury zone and epicardium, likely associated to *Axl* expression in injury zone macrophages and in *Pkd1*+ epicardial cells, respectively. As predicted by the L-R map, *Axl* expression levels were enriched in the CM.IS and CM.Prol.IS population compared to homeostatic state (Fig. 2k), while the canonical ligand, *Gas6*, was expressed by multiple cell types, however enriched in EC (*Lox*+) population (Fig. 2l). To verify that such spatial relationship indeed happens *in-vivo* we performed hybridization chain reaction (HCR) staining for *Axl* and *Gas6* and saw that both *Axl* and *Gas6* are significantly upregulated in the border zone compared to the remote zone and are accumulating over time (Fig. 2m-o). Taken together, after injury, a plethora of new cell states and cell types are found in the heart, generating distinct cellular interaction networks. The newly formed border zone CMs change their gene expression profile and engage in a new signaling network that includes the Gas6-Axl axis.

### *Axl* is required for axolotl border zone cell cycle activity

Having identified AXL to be enriched in the injury responsive CMs, we next aimed to check its requirement in regeneration. Given that receptor tyrosine kinases (RTKs) such as AXL signal through ligand-induced activation of their intracellular kinase domain, we followed the established strategy of overexpressing a truncated form lacking the kinase region to sequester relevant ligands without triggering downstream signaling, thereby generating a dominant-negative (dn) construct (*51, 52*). Furthermore, we added a Flag tag on the dnAXL to allow direct immunolocalization of dnAXL. We generated *CAGGs:loxP-EGFP-loxP-dnAXL-T2A-mCherry* transgenic animals that were crossed with CM specific x*CMLC2 (MYL7):-ER-CRE-ERT2* animals. Upon 4-Hydroxytamoxifen (4-OHT) treatment, animals showed dnAXL expression in scattered CMs that were also mCherry-positive and EGFP-negative (Fig. 3a,b). Staining against AXL by anti-Flag-tag antibody or HCR directed against AXL (Fig. 3b,c), showed that regions lacking the EGFP signal display stronger signal compared to eGFP+ cells. Measurement of EdU 7 days after cryoinjury, which according to our data marks the peak of cycling cells (Fig. S1c,d), showed a trend to reduced proliferation of dnAXL+ cells, although cycling cells could still be found (Fig. 3d). We next wanted to know if the absence of active AXL could affect the formation of the border zone CMs. For that, we stained for *Nppa* (Fig. 3e-g) and *Ankrd1* (Fig. 3h,i), two evolutionary conserved border zone markers (*40*), and found that they are expressed in a lower proportion of dnAXL+ CMs compared to ctrl (Fig. 3g,i). Given the apparent lower ability of dnAXL+ cells to transition into border zone CMs, we next tested whether dnAXL mosaic OE affects cardiac regeneration. Detection of fibrosis by Mason’s Trichrome and CM fluorescence reporting to evaluate scar size 2 month after injury showed a larger injured area in dnAXL hearts compared to control hearts (Fig. 3j-m, Fig. S10a-c). Altogether this data suggests that loss of active AXL in CMs leads to lower regenerative response associated to their lower ability to activate a border zone program eliciting de-differentiation and proliferation.

**Figure 3.**
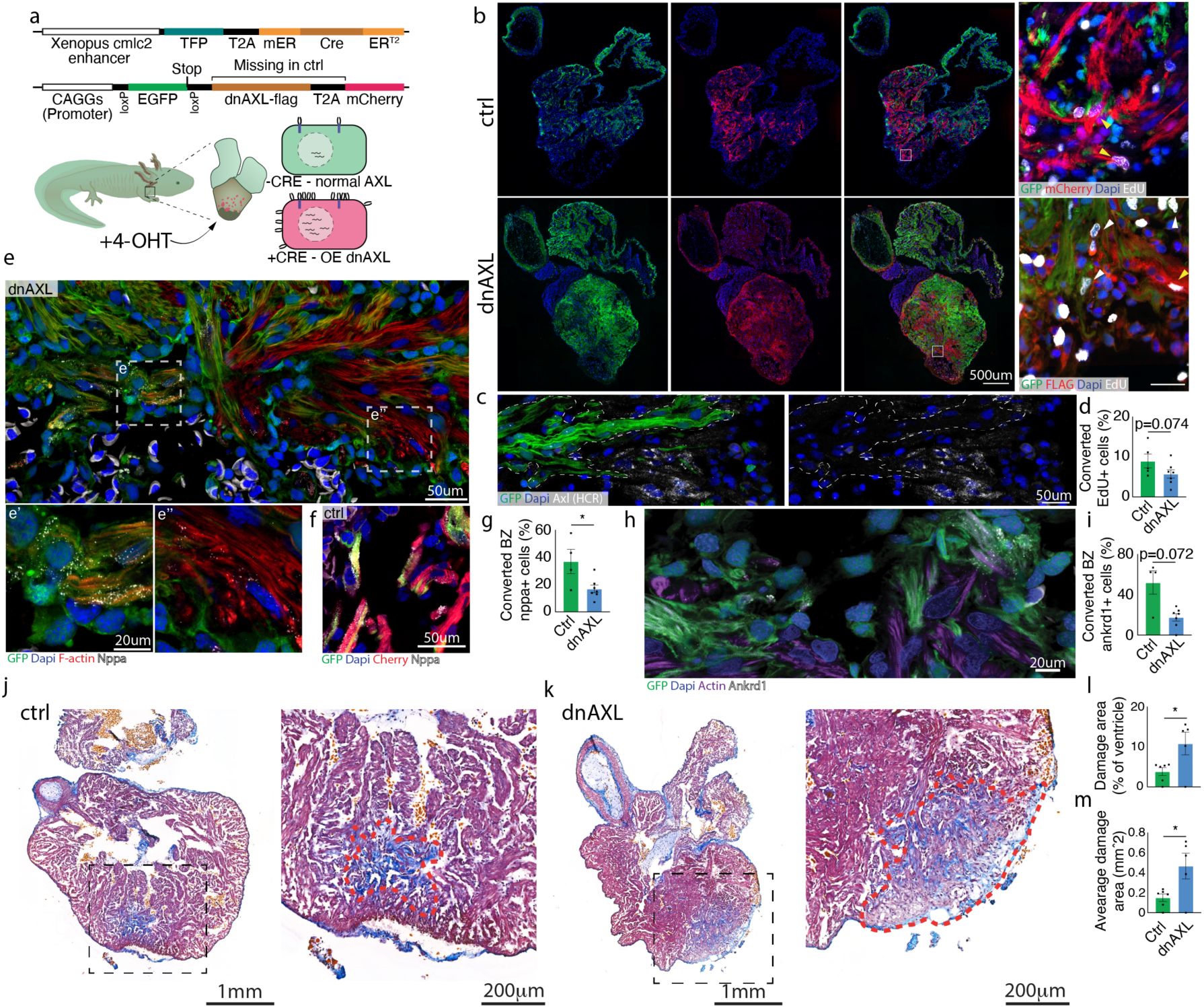
Axolotl AXL dnAXL impairs cardiac regeneration. **a**, Schematic representation of the generated transgenic animals and the expected outcome after tamoxifen induction. **b**, Representative images of ctrl and dnAXL heart sections showing mosaic pattern of endogenous GFP+ (green, non-converted), endogenous mCherry+ (red, converted in ctrl) and stained Flag-tag (red, converted in dnAXL). Squares show magnified regions stained with EdU showing cycling CMs. Yellow arrowheads show converted EdU+ CM. White arrowheads show non-converted EdU+ CM. **c**, HCR staining for Axl showing no expression in non-converted CMs while having high expression in adjacent cells. **d**, Quantification of EdU+ converted CMs. **e**-**i** HCR staining and quantification of number of positive cells for known border zone markers *Nppa* (**e**-g) and *Ankrd1* (**h,i**) of dnAXL and ctrl hearts. **j**-**m**, Mason’s trichrome staining on ctrl (**j**) and dnAXL (**k**) heart sections, 8 weeks post cryo injury showing the damage area and residual fibrotic region (pale color and blue region, marked by red dashed line). **l**,**m,** quantification of damage area as taken by the 3 sections which showed the largest damage area. Data is presented as damage area relative to size of ventricle (**l**) and average area size (**m**). \**P* < 0.05, Statistical test in all data: Mann-Whitney one-tailed T-test.

### AXL promotes dedifferentiation of mammalian CMs

After showing AXL requirement for border zone CMs in axolotl, we next asked whether AXL was sufficient to promote an effect in mammalian CMs. Previous studies have not detected AXL expression in neonatal (*53*) and adult (*22*) CMs (Fig. S11a,b), thus we devised an overexpression strategy. We generated AAV of serotype 9 (AAV9), which has a high tropism to CMs in the heart (*54*), in which either Axl or eGFP expression was under the control of a Cytomegalovirus (CMV) promoter. To determine the direct effect of AXL on CM phenotype and avoid confounding non-autonomous effects, we overexpressed AXL in primary mouse CM *in-vitro*. We isolated postnatal (P) P1 or P8 CMs specifically expressing tdTomato induced by Myh6-Cre expression and infected them with AAV9-CMV-AXL. Five days after infection, AXL OE promoted robust and reproducible sarcomere disassembly compared to non-infected cells in the same culture or AAV9-EGFP infected wells (Fig. 4a, Fig. S12a,b). Overexpression of AXL alone was sufficient to have an effect even without adding its canonical ligand, Gas6. This activation may result from the known presence of Gas6 in serum (*55*) or by self-activation in high expression conditions (*56*). As P1 cell isolations resulted in higher purity of CM and both P1 and P8 showed similar sarcomere disassembly response, further analyses were performed on P1 cells. Using a highly specific small molecule inhibitor against AXL’s catalytic kinase domain (R428) (*57*), we observed a trend to prevention of sarcomere disassembly by AXL overexpression in a dose dependent manner, suggesting that AXL signaling is required for the disassembly (Fig. 4b,c). To test the ability of AXL to influence CM sarcomere integrity *in-vivo* we injected AAV9-AXL driven by the cardiac troponin T (cTnT) promoter at P1 and harvested the hearts 8 weeks later. Using our automated sarcomere segmentation pipeline, we analyzed the a-actinin striation pattern and saw that in AXL OE cells, both area and intensity of a-actinin were lower compared to control (Fig. 4 d-h), confirming our *in-vitro* disassembly results. In many pro-regenerative overexpressing systems, such as ERBB2 (*15*) or YAP (*17*) over activation, robust sarcomere disassembly is indicative of CM dedifferentiation as well as induction of proliferation, therefore we tested whether AXL OE could induce cell cycle activity. Despite inducing sarcomere disassembly, AXL OE did not activate the cell cycle in CMs (Fig. 4i). To assess whether sarcomere disassembly is indicative of dedifferentiation, we performed bulk-RNAseq on day 3 after infection of AXL/GFP. AXL OE dramatically changed the transcriptome with ±7711 differently expressed genes (adjusted P value<0.05) (Fig. 4j, Fig. S13a). On the top of the DEGs list, several known border zone CMs genes (*40*) such as *Clu*, *Nppa* and *Nppb* were upregulated (Fig. 4j). Looking at a comprehensive list of border zone transcriptomic markers, showed that out of the 9 known genes, 7 were significantly upregulated in AXL vs GFP samples, whereas the remote zone markers were higher in the GFP cells (Fig. 4k). To broadly understand the effects of AXL OE on the cells, we used ingenuity pathway analysis (IPA), which showed enrichment in cancer and development related ontologies, including “invasion of cells” and “migration of cells” also characteristic of regenerating tissues (Fig. S13b). Among the most inhibited ontologies, we observed cell death related terms such as “apoptosis of lymphatic system cells” and “organismal death” (Fig. S10b). To assess whether AXL can reduce CM apoptosis, we treated isolated neonatal CMs with AAV9-AXL or AAV9-GFP and exposed them to either normoxic (21% O^2^) or hypoxic (3% O_2_) conditions. Under normoxia, both groups exhibited similar levels of apoptosis, as indicated by Terminal Deoxynucleotidyl Transferase dUTP Nick End Labeling assay (TUNEL) positive cells. However, under hypoxia, AXL-treated cells-maintained baseline levels of apoptosis, whereas GFP-treated cells showed threefold increase in cell death. Thus, AXL likely rescues CMs from hypoxic cell death compared to the ctrl (Fig. S13c).

**Figure 4.**
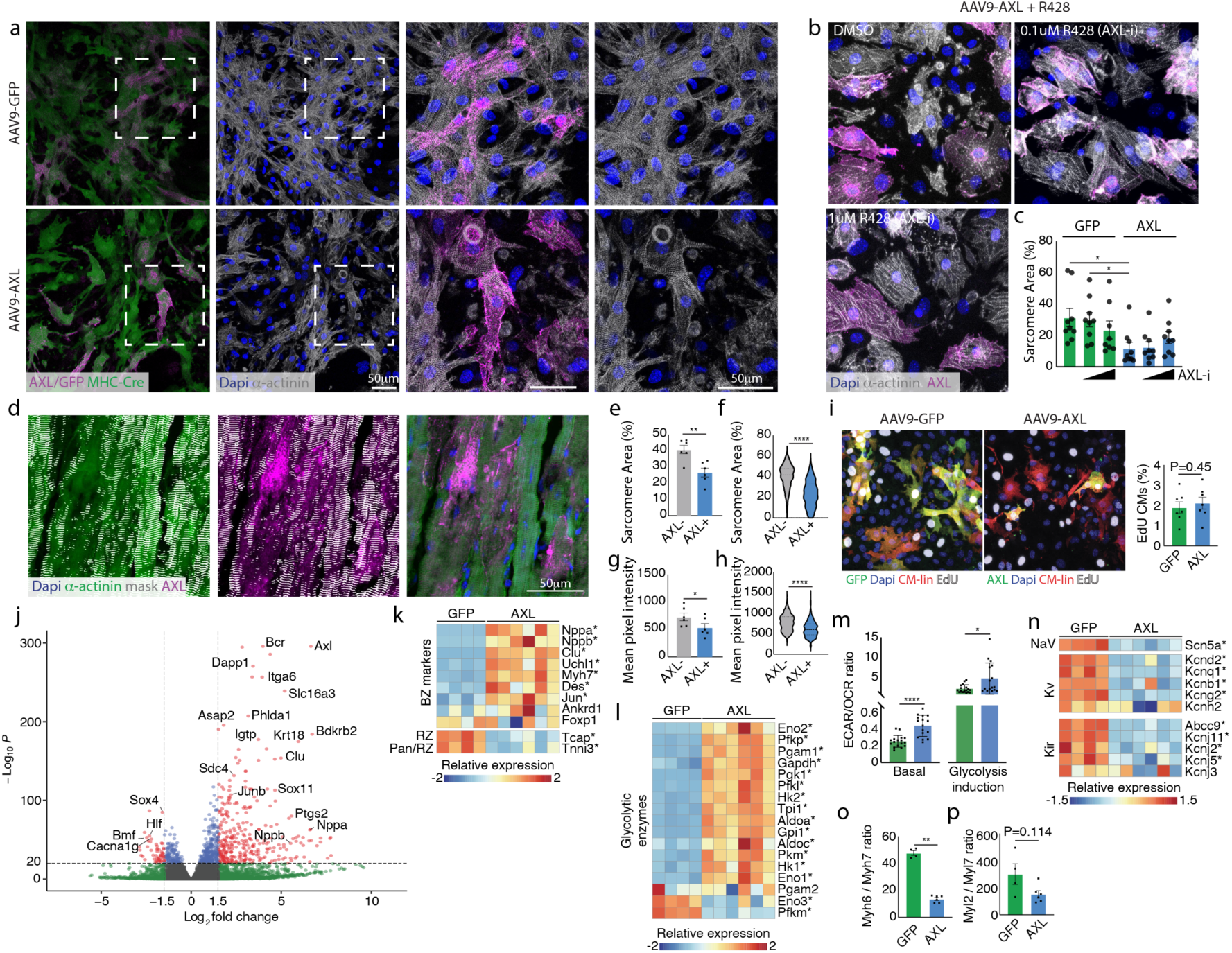
Overexpression of axl in primary mouse cardiomyocytes. **a**, Representative images of P1 neonatal CMs 5 days after AAV9-AXL or AAV9-GFP infection stained for AXL or GFP (magenta), linage labeling of CMs (green), α-actinin (gray) and dapi. **b**, Representative images same as (**a**) only with addition of 0uM, 0.1uM or 1uM R428 (axl inhibitor). **c**, Quantification of sarcomere integrity from **a,b. d-h**, *in-vivo* AAV9-AXL/GFP administration at neonatal P1 and analyzed at 8 weeks. Representative image of AXL induced sarcomere disassembly (**d**) as evident by less α-actinin staining as calculated by automated sarcomere detection per area (average between samples in **e**, all cells combined in **f**) and lower mean α-actinin mean pixel intensity (average between samples in **g**, all cells combined in **h**). n=504 AXL-cells and 709 AXL+ cells. **i,** Representative images and quantification of EdU cell cycle indicators of P1 CMs *in-vitro,* 3 days after addition of AAV9-AXL or AAV9-GFP. **j**, Volcano plot generated from bulk-RNA seq of 6xAAV9-AXL and 4xAAV9-GFP samples, of P1 CMs *in-vitro*, 3 days after infection. Genes in red are significantly up or downregulated (1.5 > log2FC, -1.5 < log2FC, -log10P < 1xe-20), as calculated by DESeq2. Gene in blue are significant but not sufficiently changed. Genes in green are sufficiently up or down regulated but not significantly. **k**, Heatmap of known border zone or remote zone markers in AAV9-AXL (AXL) or AAV9-GFP (GFP) taken from bulk RNA-seq. Asterisk next to gene name shows p<0.05 using DESeq2 statistical test. **m**, Heatmap of known glycolytic enzymes in AAV9-AXL (AXL) or AAV9-GFP (GFP) taken from bulk RNA-seq. Asterisk next to gene name shows p<0.05 using DESeq2 statistical test. **m**, Ratio of Extracellular acidification rate (ECAR)/ Oxygen consumption rate (OCR) indicating the preference of pyruvate fate of either fermentation or oxidation by mitochondria. Glycolysis induction was calculated by subtraction of values after 2-Deoxy-D-glucose induction from glucose induction values. **n**, Heatmap of major ion channels selected from expression levels of adult mouse hearts in AAV9-AXL (AXL) or AAV9-GFP (GFP) taken from bulk RNA-seq. Asterisk next to gene name shows p<0.05 using DESeq2 statistical test. **o**, Myh6/Myh7 gene expression ratio as calculated from bulk RNA-seq normalized counts. **p**, Myl2/Myl7 gene expression ratio as calculated from bulk RNA-seq normalized counts. In all relevant panels \**P* < 0.05, \*\***P** < 0.01, \*\*\***P** < 0.001 ****P<0.0001 (statistical test: Mann-Whitney two-tailed T-test). In panels k,l and n, asterisk shows significant change as calculated by DESeq2.

To gain further insights into the cell-state changes induced by AXL expression in CMs we used Kyoto Encyclopedia of Genes and Genomes (KEGG) database to look for processes enriched in our dataset. Half of the top 10 most enriched terms were related to cell metabolism, including terms like: “oxidative phosphorylation”, “Fatty Acid Metabolism” and “Citrate Cycle TCA Cycle” (Fig. S13d). As reduction in Glycolysis derive pyruvate to Ox-Phos and concomitant increase in lactic fermentation of pyruvate are additional hallmarks of dedifferentiation (*2*), we next focused on changes occurring in specific groups of metabolic genes. Looking specifically at glycolytic enzyme genes, a significant increase in 14 out of the 17 genes could be seen (Fig. 4h). A similar trend could also be seen by looking at a general glycolysis gene list (Fig. S13a). In sharp contrast, genes encoding structural subunits of OxPhos mitochondrial complexes I (95%), II (100%), III (90%) and V (90%) were downregulated in the AXL treated cells (Fig. S14b,c). Genes encoding proteins of complex IV (CIV) showed a complex pattern coherent with dedifferentiation (Fig. S14b). Unlike other complexes, 7 out the 14 structural subunits of CIV (*Cox4l*, *Cox6a*, *Cox6b*, *Cox7a*; *Cox7b*, *Cox8* and *Ndufa4*) have paralogs that are expressed upon metabolic switch (*Cox4l2*, *Naduf4al2*) (*58*) or upon cardiomyocyte maturation (*Cox6b*, *Cox6a2*, *Cox7a1*) (*58, 59*). Of interest, overexpression of AXL promotes the switch of CIV paralog genes towards a more hypoxic profile (switch from *Cox4l1* to *Cox4l2* and *Ndufa4* to *Ndufa4l2*) and towards the non-heart specific paralogs (*Cox6a1*, *Cox8a*, *Cox7a2*) (Fig.S14b). Such specific switch in CIV paralogs indicated the fine-tuning capacity of AXL for triggering the dedifferentiation. In addition, genes encoding fatty acid oxidation enzymes showed a marked decrease in AXL OE group (Fig. S14c). To functionally assess metabolic state, oxygen consumption rate (OCR) and extracellular acidification rate (ECAR) were measured using Seahorse assays. Higher ECAR/OCR ratio implies a preferential use of pyruvate toward lactate fermentation, whereas lower ratio corresponds to preferential routing toward mitochondrial oxidation. AXL OE displayed a higher basal ECAR/OCR ratio (Fig. 4m), which became significantly elevated after stimulation of glycolysis in the Glycolysis Stress assay (Fig. 4m). In the Cell Mito Stress assay, AXL OE, also showed a significantly increased ECAR/OCR ratio under coupled respiration (Fig. S14d) which was abrogated and similar in both groups following FCCP-mediated uncoupling respiration, when mitochondrial capacity is saturated.

Altogether these results show that AXL expression modify the CM metabolic program driving pyruvate fate towards to fermentation at the expenses of oxidation by mitochondria and validating our RNA-seq results.

We also examined whether AXL-expressing CMs show differences in CM maturation markers compared to controls, such as levels of K+ and Na+ channel expression, which regulate membrane potential and electrical impulse. Focusing on the top 5 channels expressed in the adult mouse heart from each type (taken from a published dataset (*60*)), we first focused on components of the inwardly rectifying potassium (Kir) channels. All top-5 expressed genes of this family (*Abcc9*, *Kcnj2/3/5/11*) were downregulated in the AXL OE vs GFP CMs (Fig. 4o). Similar downregulation was observed within the Voltage-gated potassium (Kv) channels (*Kcnd2/q1/b1/g2*) with the exception of Kcnh2, which did not show any different levels between groups (Fig. 4o). Lastly, the main cardiac sodium channel Scn5a (Nav1.5) is dramatically downregulated in AXL-expressing cells, altogether suggesting that AXL promotes an immature CM ion channel expression profile. To further assess of dedifferentiation, we examined the expression pattern of two well defined parameters: the ratios of *Mhy6/Myh7* and *Myl2/Myl7*, both indicative of the “maturity” levels of the CMs. In both gene assays, the GFP treated group showed higher expression ratios, further suggesting less mature state in AXL OE CMs (Fig. 4p,q). Unsurprisingly, given the lack of added proliferation after AXL treatment, genes related to cell cycle activity showed no obvious trend except for DNA replication, licensing and initiation genes which were higher in the GFP group (Fig. S15). Altogether, AXL overexpression in mammalian CMs promotes robust dedifferentiation as seen by several well characterized dedifferentiation hallmarks, while not promoting cell cycle activity.

## Discussion

One of the central promises of regenerative biology is the discovery of novel therapeutic targets by studying species with high regenerative capacity. To this end, we generated the first comprehensive spatial multi-omics atlas of axolotl cardiac regeneration, enabling the identification of cell-type-specific markers across all major cardiac populations. This high-resolution resource allowed us to resolve an injury-specific CM trajectory and identify possible regulators of the regenerative response. Among these, *Axl* was robustly upregulated in border zone CMs, with its canonical ligand, *Gas6*, primarily expressed by endothelial cells, placing it among a growing group of angiocrine factors potentially involved in cardiac regeneration.

AXL belongs to the TAM (TYRO3, AXL, MERTK) receptor family, which is broadly involved in immunomodulation, including apoptotic cell clearance and resolution of inflammation. In the context of mammalian heart injury, AXL is enriched in pro-inflammatory macrophages and was shown to promote adverse cardiac effects in ischemia/reperfusion infarction injuries (*61*). The suggested mechanism through which AXL promoted the macrophage inflammatory profile was acidification of the cytoplasm and induction of a glycolytic metabolic shift (*61*). Although these metabolic shift in macrophages is detrimental for the injured heart, a similar metabolic shift in CMs, as observed here, is reminiscent of an embryonic metabolic profile, a hallmark of CM dedifferentiation and protective against hypoxic shock in CMs.

In addition to metabolic shift, AXL overexpression induced robust sarcomere disassembly and altered expression of ion channels, yet it failed to initiate CM proliferation. These findings indicate that dedifferentiation can be uncoupled from cell cycle re-entry, challenging the assumption that these processes are necessarily linked. Our data support a growing body of evidence indicating that certain dedifferentiation features, such as sarcomere disassembly, are not sufficient to induce proliferation (*62*). These insights raise the question of whether cell cycle re-entry should remain a core criterion in the definition of CM dedifferentiation. It is also possible that the AXL-OE brings CMs to a non-mitogenic but primed state that could be more responsive to additional, mildly pro-proliferative cues such as Agrin or Periostin. This suggests that dedifferentiation may serve not as a direct trigger, but as a permissive state that enhances CM plasticity and regenerative competence in response to subsequent stimuli.

In line with this, AXL has been implicated in HER2+ breast cancers, where it heterodimerizes with ERBB2, a well-established inducer of cardiac regeneration, to promote metastasis, but is dispensable for primary tumor growth, suggesting that ERBB2 and AXL have non-redundant roles in tumor progression and that AXL may not serve as a mitogenic signal on its own (*43*). Published mouse datasets revealed that *Axl* is absent in CMs during both neonatal (*53*) and adult (*22*) stages (Fig. S11a,b). Considering the strong effect of constitutively active ERBB2 on CM proliferation and dedifferentiation an intriguing possibility is that ERBB2 works in part by upregulating AXL.

Further supporting its dedifferentiation activity, AXL has been identified as a transcriptional target of the AP-1 transcription factor family (*63*), which has emerged as a key regulator of regeneration across multiple species and tissues (*8, 40, 64*). AP-1 factors are known to regulate chromatin accessibility and promote cytoskeletal remodeling, including sarcomere disassembly and cellular protrusion during zebrafish heart regeneration (*8*). Given our observation of robust dedifferentiation following AXL overexpression, it is tempting to speculate that AXL may mediate a subset of AP-1–induced regenerative effects. Moreover, as AP-1 mediated responses are conserved across diverse regenerative contexts, further exploration into the breadth of AXL’s involvement could help clarify its role as a generalized regulator of dedifferentiation.

In this study, we identified AXL to be upregulated in regenerating CMs and as a driver of dedifferentiation, we speculate that AP-1 TF family upregulation in BZ-CMs upregulates its expression, which leads to metabolic and structural changes involved in regeneration and that may render CMs susceptible to additional mitogenic cues. Furthermore, here we focused on the cell-autonomous effects of AXL in mammalian CMs, allowing us to dissect dedifferentiation from cell cycle activation pathways. The *in-vivo* consequences of AXL activation, particularly in the context of the heart’s exaggerated inflammatory response, remain an important avenue for future exploration.

## Supporting information

Video 1 - uninjured

Video 2 - 14d PCI

Video 3 - 28d PCI

## Acknowledgements

We thank the Tanaka and Torres laboratories for their discussions, input, and support. During editing of this manuscript, Chat-GPT was used for text style improvement. We thank Histology, NGS and Bio-optics from the Vienna BioCenter Core Facilities (VBCF) and Vienna BioCenter animal caretaker team for continuous outstanding support and services. We thank members of the Microscopy and Dynamic Imaging, Viral Vectors, and Animal Facility CNIC units for excellent support. We thank Eldad Tzahor, Nadia Mercader and all other members of the REANIMA consortium as well as Kenneth Poss for fruitful discussions and shaping this work. We thank the labs of Rubén Marín-Juez, Kristy Red-Horse and Karina Yaniv for help with cell type annotation.

## Author contribution

Designed study: E.B., J.P.P., M.T., and E.M.T. Performed bioinformatic analysis: J.W., I.R.G., F.J.F.C., F.S.C., T.L. and E.B. Generated transgenic axolotls: Y.T.S., E.B., K.L., and J.Y. Prepared and collected axolotl samples: E.B., and A.P. Prepared and collected mouse samples: J.P.P., L.D., A.P., and E.B. Performed and analyzed metabolic study: J.P.P., P.F.M., and J.A.E. Prepared the figures, wrote and approved the manuscript: E.B., J.W., J.P.P., M.T., and E.M.T.

---

**Figure S1.**
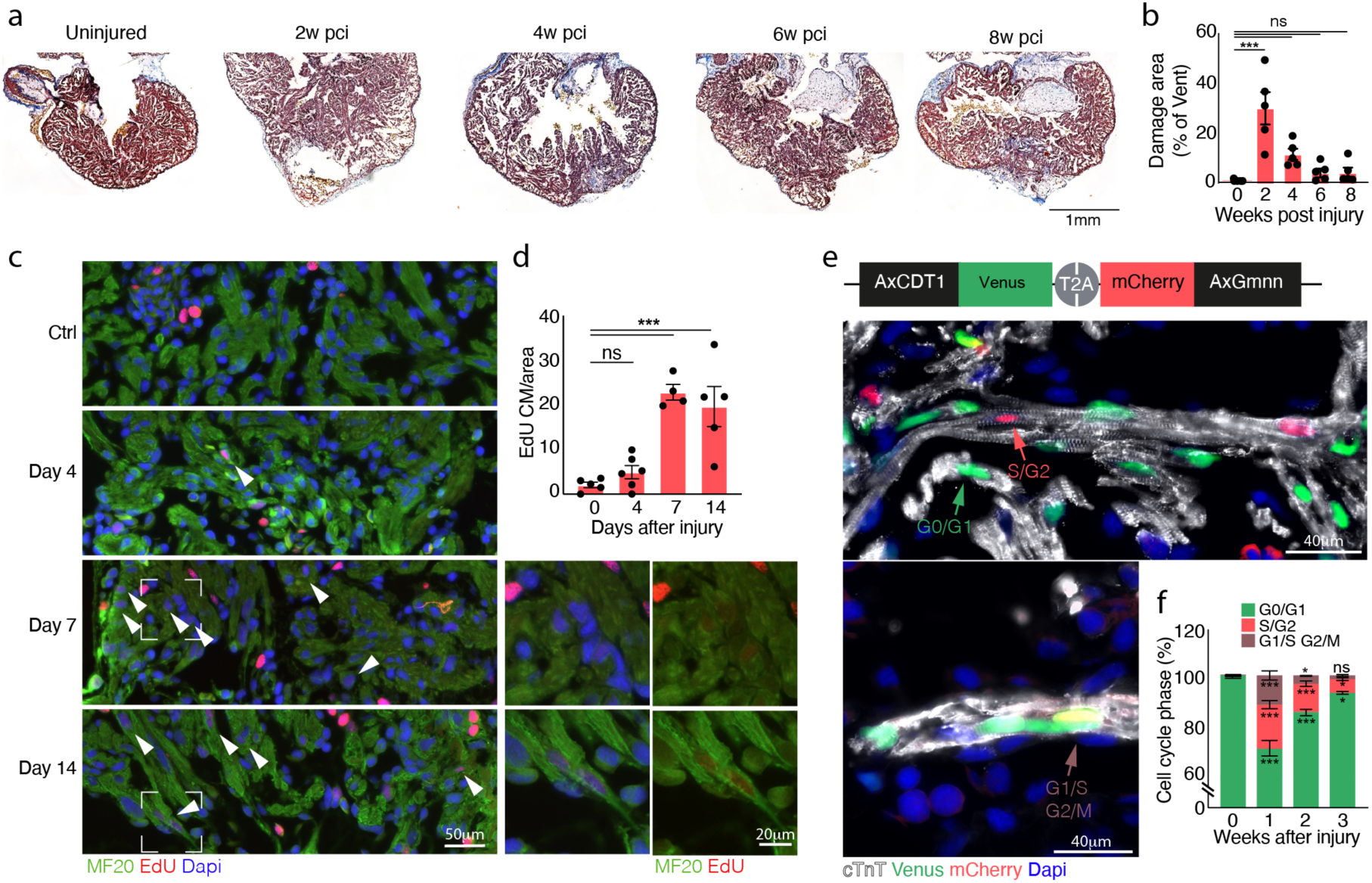
Characterization of axolotl cardiac injury. **a**, Representative images of cardiac sections stained with mason’s trichrome of time course following cardiac injury in axolotls, examined at 0, 2, 4, 6, and 8 weeks post-injury. **b**, Quantification of the damaged area over time. **c**, Representative images showing EdU incorporation at 0, 4, 7, and 14 days post-injury, indicating proliferative activity. **d**, Quantification of EdU+ cells in the injured myocardium. **e**, FUCCI-reporter heart section showing cell-cycle stage distribution: green (G0/G1), red (S/G2), yellow (G1/S and G2/M). **f**, Quantification of FUCCI+ CMs at 0, 1, 2, and 3 weeks post-injury n= 4 uninjured, 3×1 week, 3×2 weeks, 3×3weeks after injury). Data are presented as average ± s.e.m. \**P* < 0.05, \*\***P** < 0.01, \*\*\***P** < 0.001 (statistical test: two-way ANOVA with Dunnett’s post hoc, all groups were compared to control group. In **d**, Mann-Whitney two-tailed T-test).

**Figure S2.**
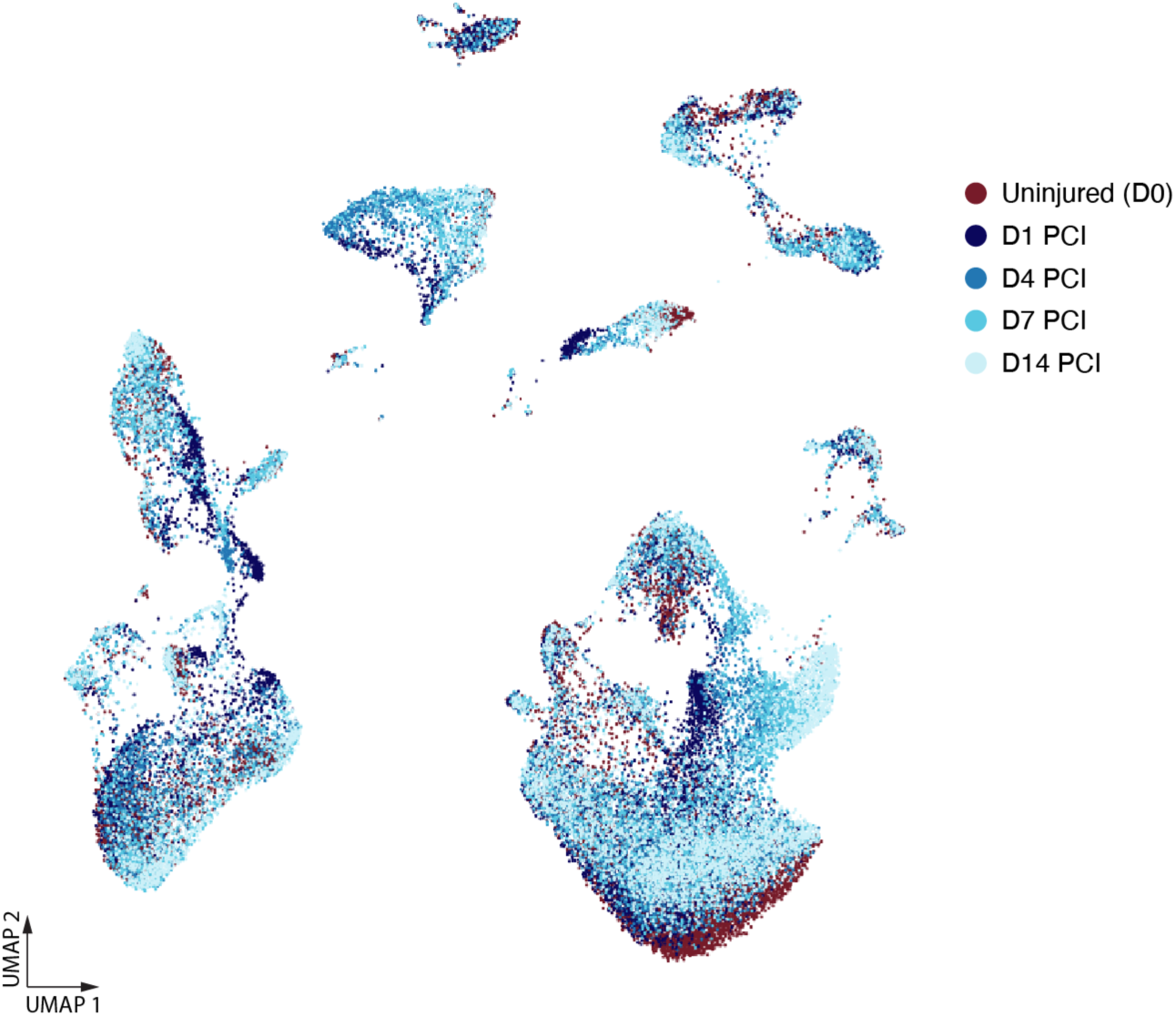
Cellular heterogeneity in the axolotl heart. UMAP depicting all the cells acquired in the snRNA-seq labeled by their relevant timepoint. Each timepoint represents cells collected from 6 polled hearts.

**Figure S3.**
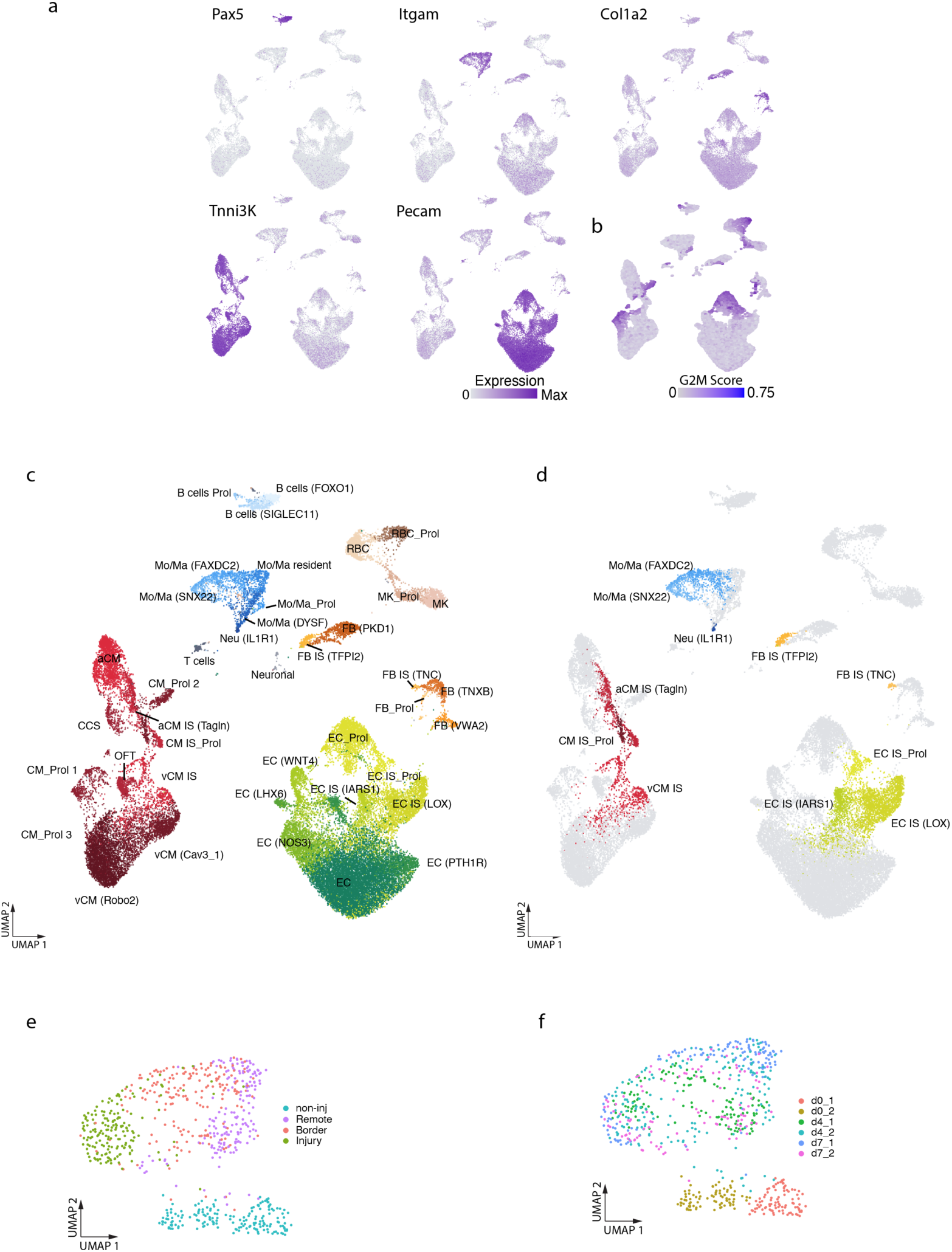
Characterization of cardiac cell populations and injury-specific clusters. **a**, Feature plots showing key markers used to identify main cardiac populations. **b**, G2M transcriptomic score depicting cell cycling cells. **c**,**d** Higer information UMAP (compared to Fig. 2a) showing all 41 annotated clusters identified in our dataset (**c**) or highlighting only injury specific clusters (**d**). **e**,**f**, UMAP depicting the clustering of individual visium “pixels” between two biological repeats of day 0,4 and 7 showing clustering by manually curated region (**e**) or by sample ID (**f**).

**Figure S4.**
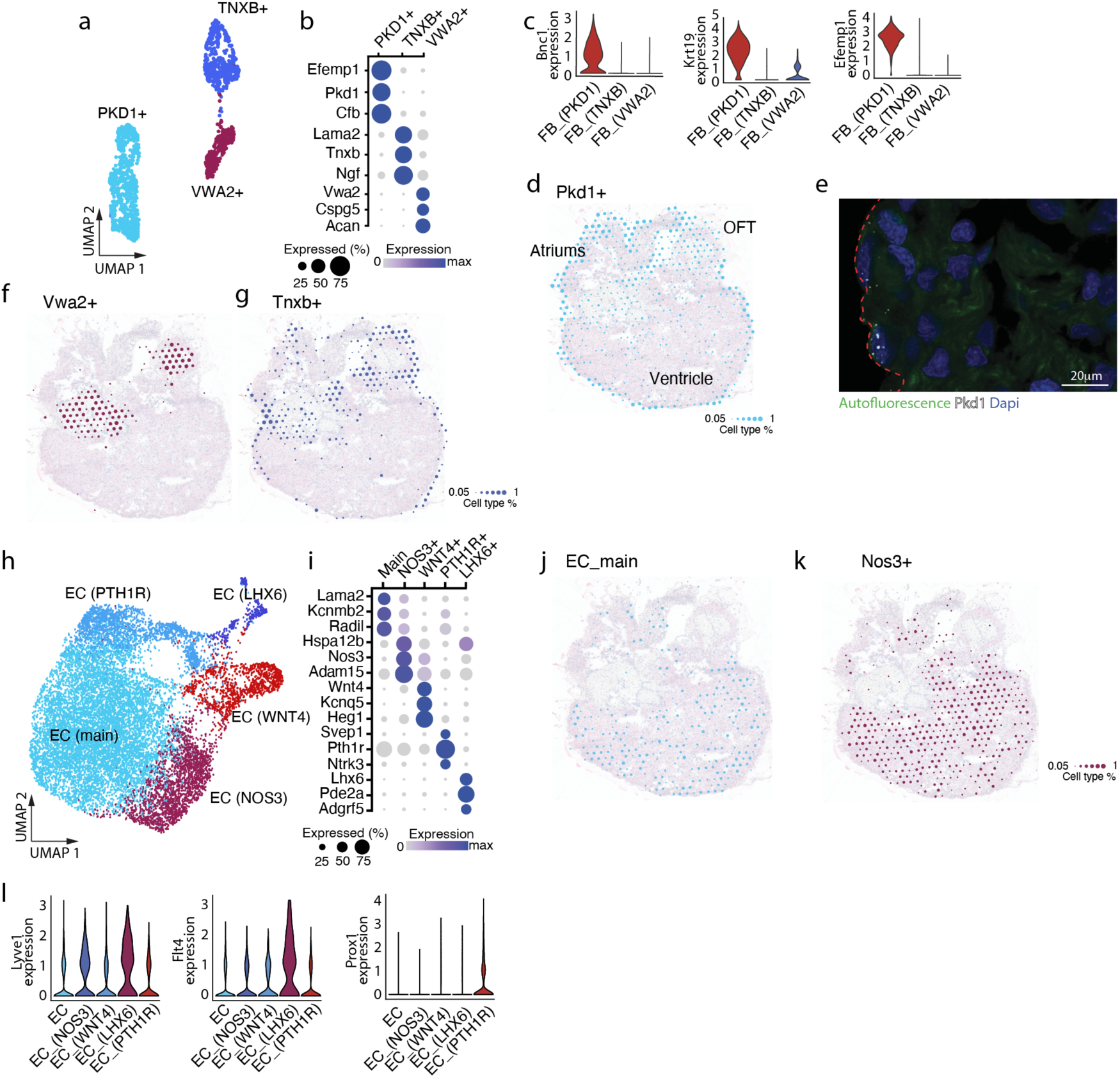
Cellular diversity of endothelial and fibroblast/epicardial cells. **a**, UMAP depicting homeostatic fibroblast populations. **b**, Dot plot showing main marker genes for each population. **c**, Violin plots showing key epicardial marker expression in homeostatic FB clusters. d, spatial mapping of *Pkd1*+ population onto the Visium slide showing mostly epicardial localization. **e**, Representative image of a non-injured axolotl heart section zooming on the edge of the heart (marked by red dashed line). Pkd1+ (white HCR staining) cells were found directly adjacent to the edge, i.e. epicardial region. Green channel marks autofluorescence to see heart contour. **f**,**g** spatial localization of *Vwa2*+ (**f**) and *Tnxb*+ (**g**) populations onto the Visium slide showing unique localization patterns in the heart. **h,** UMAP depicting homeostatic fibroblast populations. **i**, Dot plot showing main marker genes for each population. **j,k** Spatial localization of main EC (**i**) and the *Nos3*+ populations showed localization throughout the ventricle in a non-specific pattern. **l**, Violin plots showing key lymphatic marker expression in homeostatic EC clusters. In (**b**) and (**i**) dot size represent the percent of cells within each cluster expressing the gene while color shows the extent of expression.

**Figure S5.**
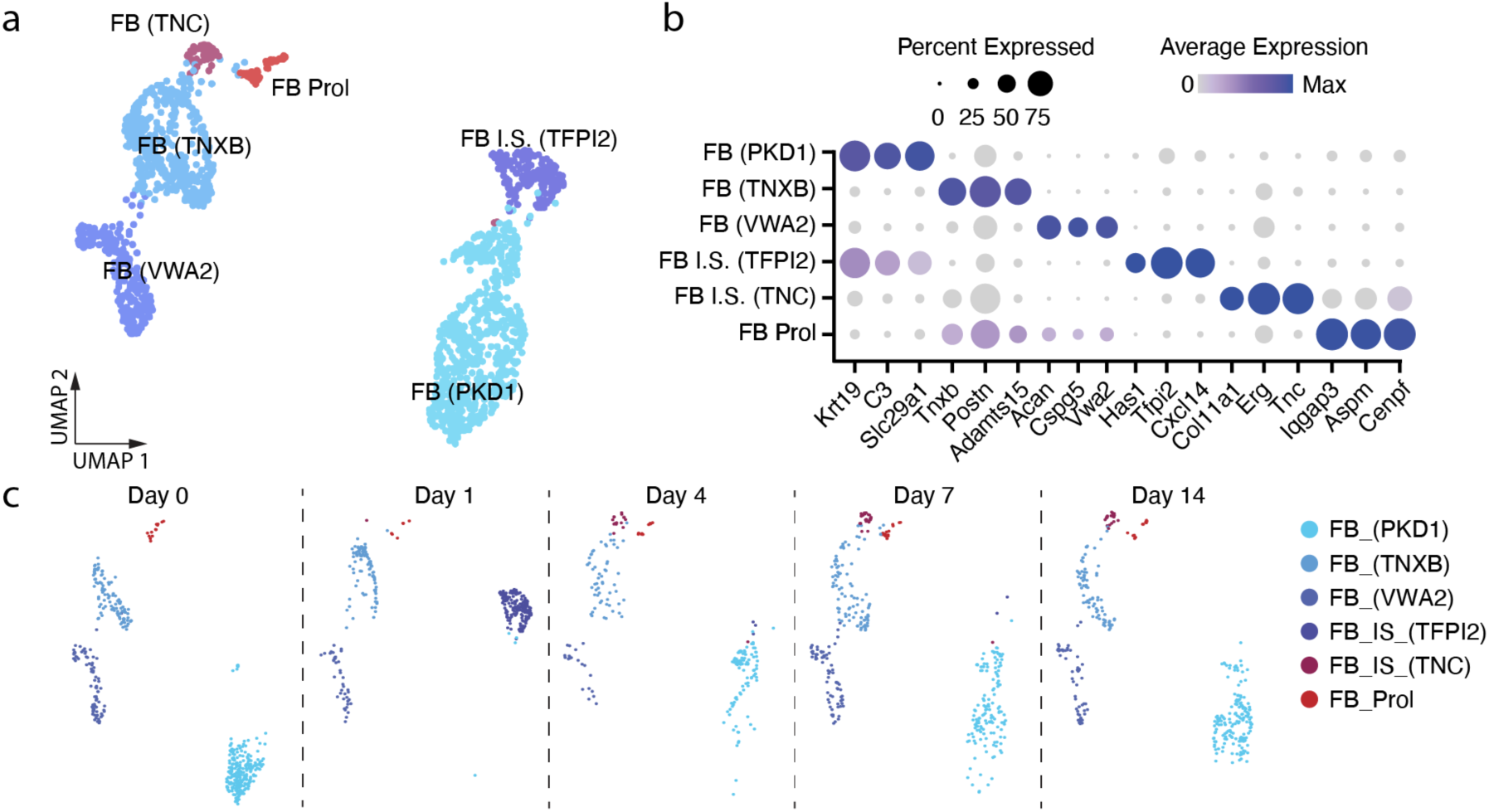
Cellular diversity of fibroblast/epicardial cells after injury. **a**, UMAP depicting fibroblast population after injury. **b**, Dot plot showing main marker genes for each population. Dot size represent the percent of cells within each cluster expressing the gene while color shows the extent of expression. **c,** UMAP showing the dynamics of fibroblast subtypes at 0,1,4,7 and 14 days post injury.

**Figure S6.**
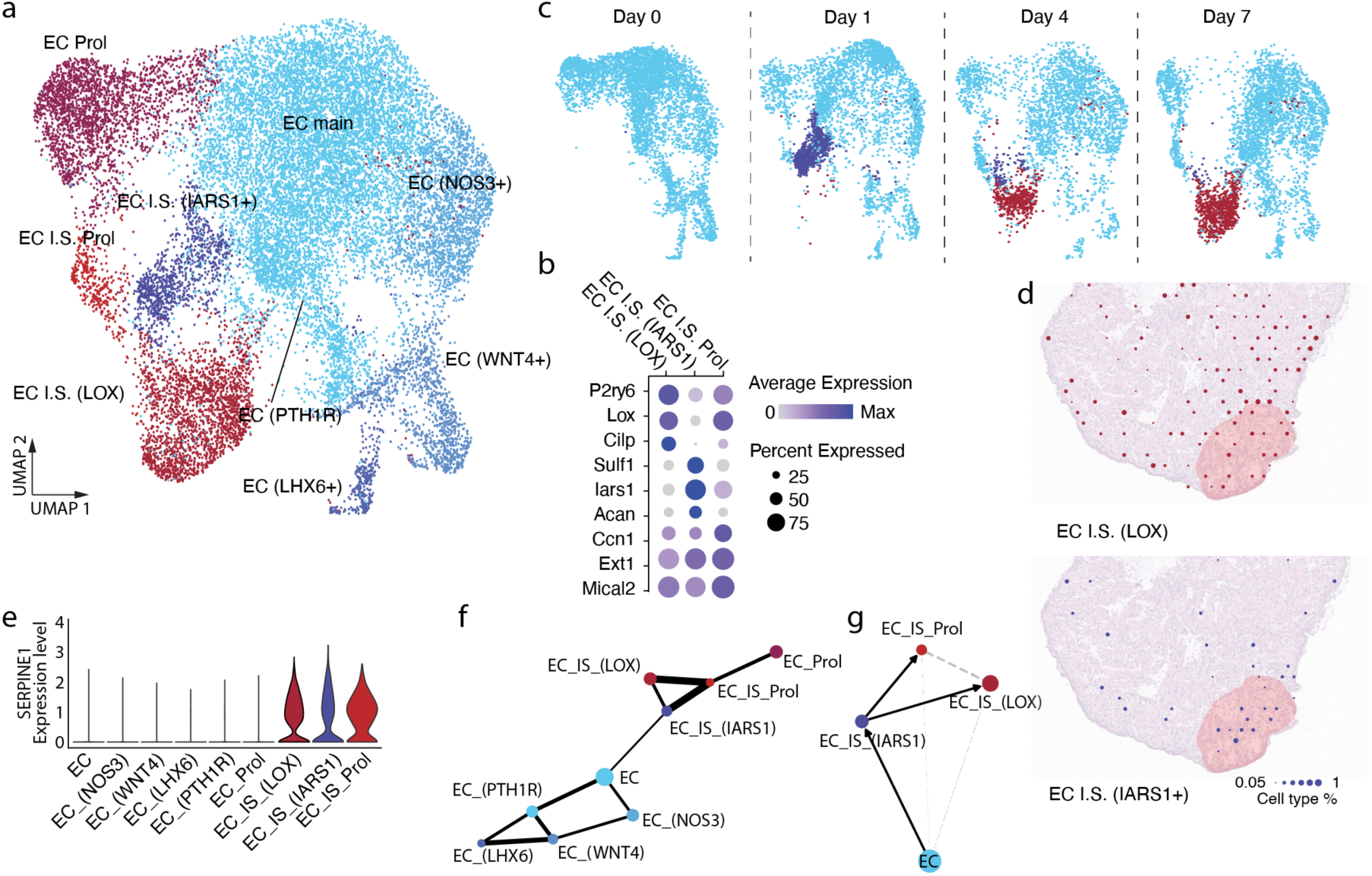
Cellular diversity of endothelial cells after injury. **a**, UMAP depicting endothelial population after injury. **b**, Dot plot showing main marker genes for each population. Dot size represents the percent of cells within each cluster expressing the gene while color shows the extent of expression. **c**, UMAP showing the dynamics of EC subtypes at 0,1,4, and 7 days post injury. Injury specific population are marked in dark blue and red colors while homeostatic populations are marked in light blue. **d**, Spatial localization of both injury specific endothelial cell populations overlayed on day 4 heart sections stained with H&E. Dots represent each Visium “pixel”. Size represents the proportion of the specific assayed cell type within the pixel. **e**, Violin plot showing the endocardial response gene, *Serpine1* expression in EC clusters. **f,g,** Partition based graph abstraction (PAGA) trajectory analysis of all endothelial subsets (**f**) and injury responsive (**g**) populations

**Figure S7.**
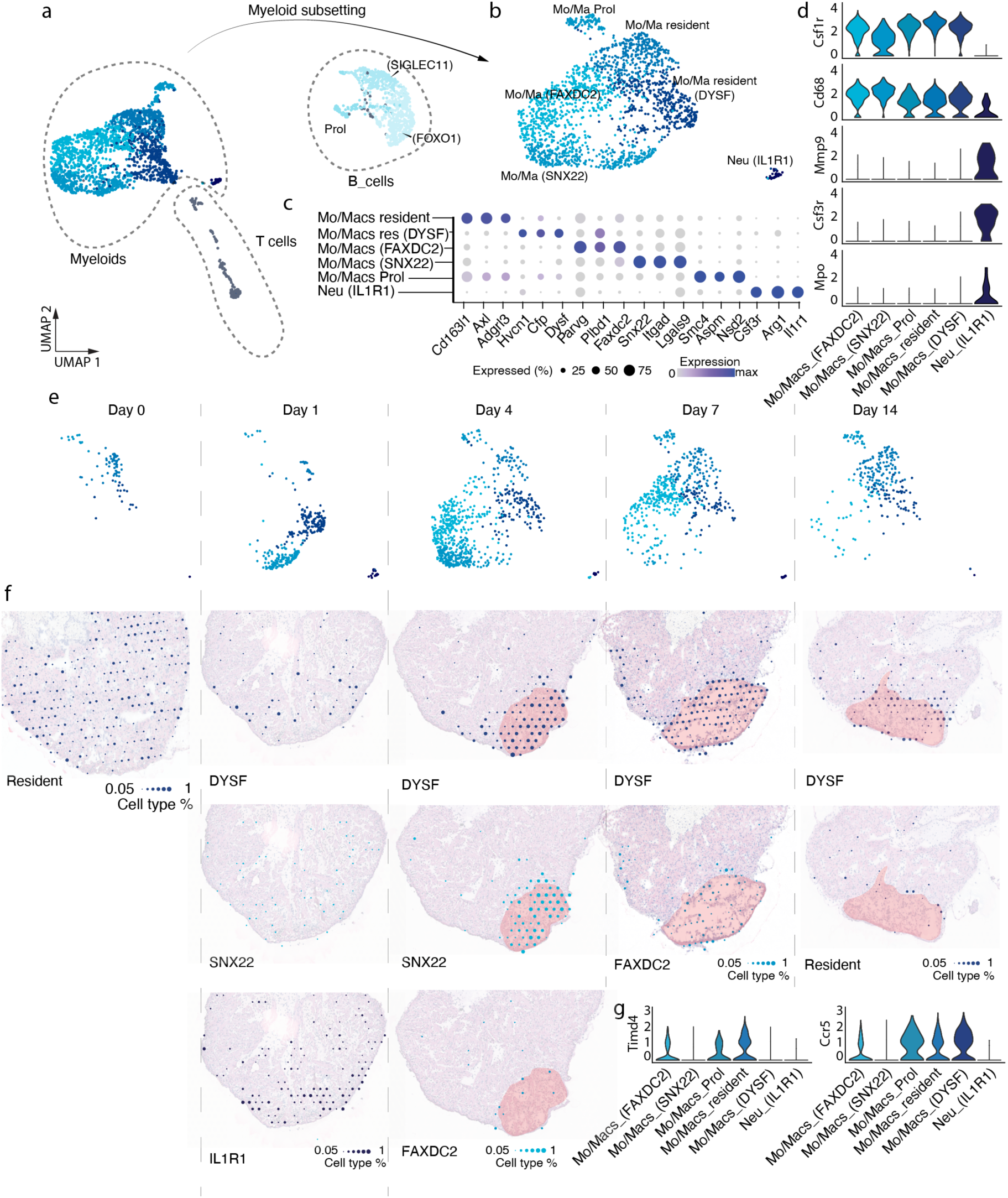
Cellular diversity and spatiotemporal dynamics of immune cells following cardiac injury. **a,b**, UMAP projections of single-cell transcriptomes showing immune cell populations (**a**) and a focused view of the myeloid cell subset (**b**) at various time points post-injury. **c**, Dot plot displaying canonical marker genes for each immune population. Dot size reflects the percentage of cells expressing the gene within each cluster; color intensity represents average expression level. **d**, Violin plots showing expression of macrophage-associated genes (Csf1r, Cd68) and neutrophil-associated genes (Mmp9, Mpo, Csf3r) across myeloid subtypes. **e**, UMAPs colored by time point, showing dynamic shifts in myeloid cell subtype composition at 0, 1, 4, 7, and 14 days post-injury. **f**, Spatial localization of major myeloid populations overlaid on H&E-stained heart sections from matching time points, illustrating infiltration and positioning in tissue context. **g**, Violin plots depicting differential expression of Timd4 and Ccr5 across myeloid clusters.

**Figure S8.**
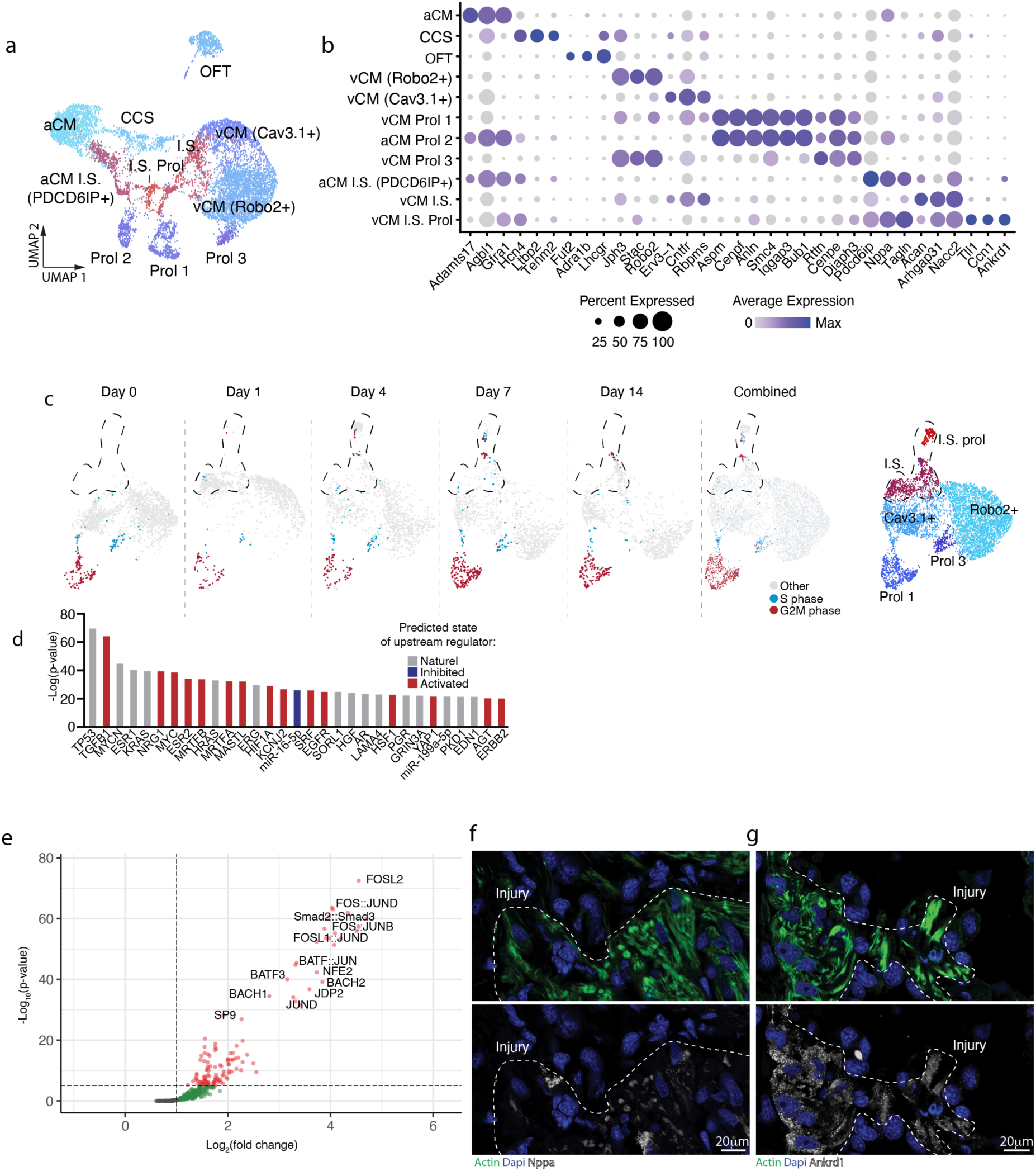
Cellular diversity of cardiomyocytes following cardiac injury. **a**, UMAP visualization of all CM populations revealing transcriptionally distinct clusters. **b**, Dot plot of canonical and injury-associated marker genes for each CM population. Dot size represents the percentage of cells within a cluster expressing the gene; color intensity indicates average expression level. **c**, Cell cycle transcriptomic score for S and G2/M phases over time depicted on the ventricular CM UMAP (depicted on the right and taken from Fig. 2a). **d**, IPA upstream regulator analysis of I.S. prol population vs the homeostatic CM populations. Colors represent predicative direction of regulator, red = activated, blue = inhabited, gray = unclear. **e,** Volcano plot showing the most enriched TF motifs in border zone CM compared to homeostatic CM populations. **f,g**, Representative HCR staining of conserved border zone marker genes *Ankrd1* (**f**) and *Nppa* (**g**) in heart sections at 4 days post-cryo injury. Dashed lines delineate the injury-core and border zone interface.

**Figure S9.**
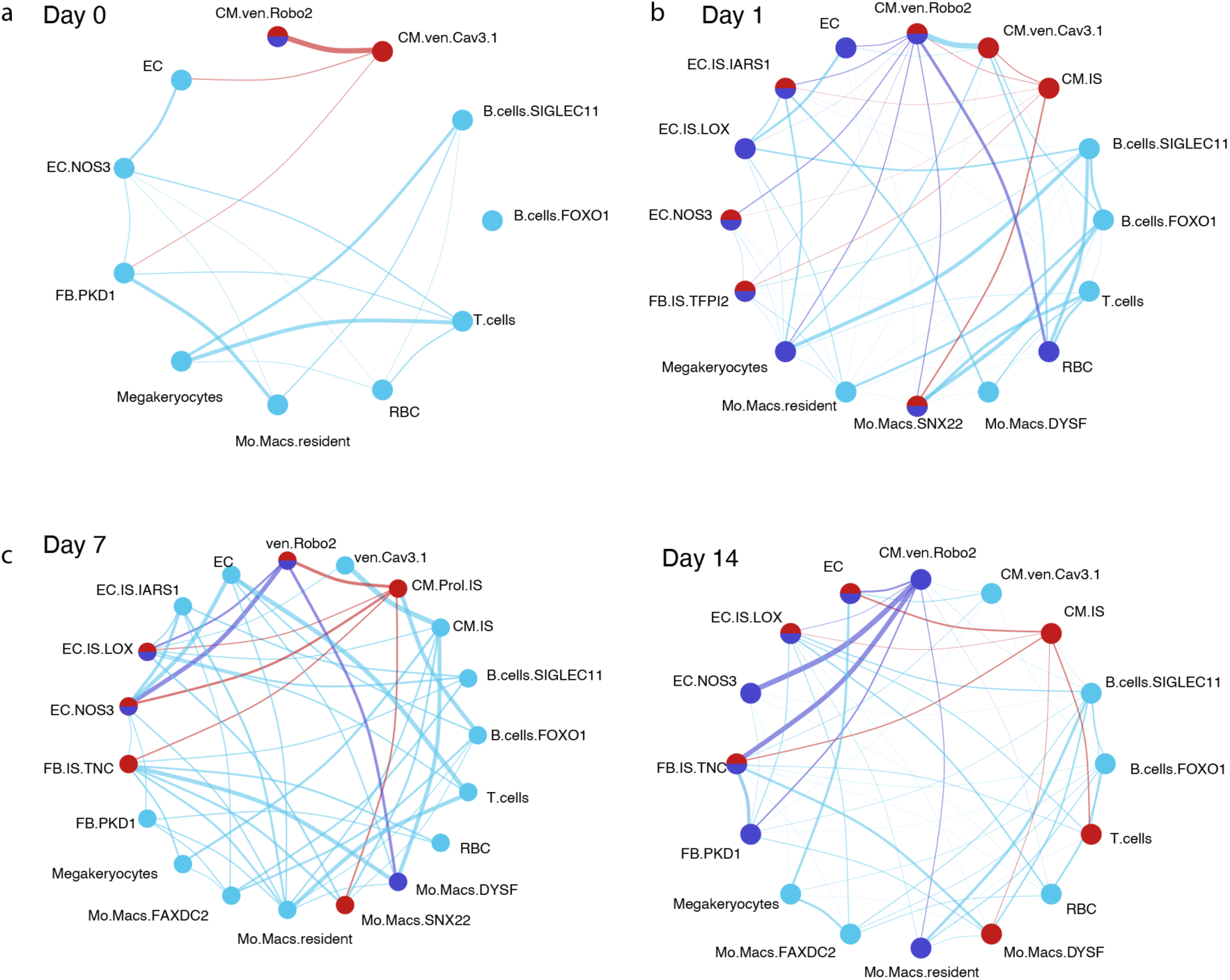
Neighborhood analysis of cardiac niche cell interactions following injury. **a–d**, Spatially resolved niche cell–cell interactions inferred using MISTy, depicting intra- and inter-pixel neighborhood interactions at day 0 (**a**), day 1 (**b**), day 7 (**c**) and day 14 (**d**) post-injury. Colors represent distinct cell type interactions; line thickness reflects interaction strength. Red nodes and lines represent the cell clusters with higher score of localizing with the CM.Prol.IS (border zone). Dark blue nodes and lines represent the cell clusters with higher score of localizing with the homeostatic Robo2+ population. Half red half blue nodes are shared between the BZ and homeostatic cells. Thicker lines represent higher localization score.

**Figure S10.**
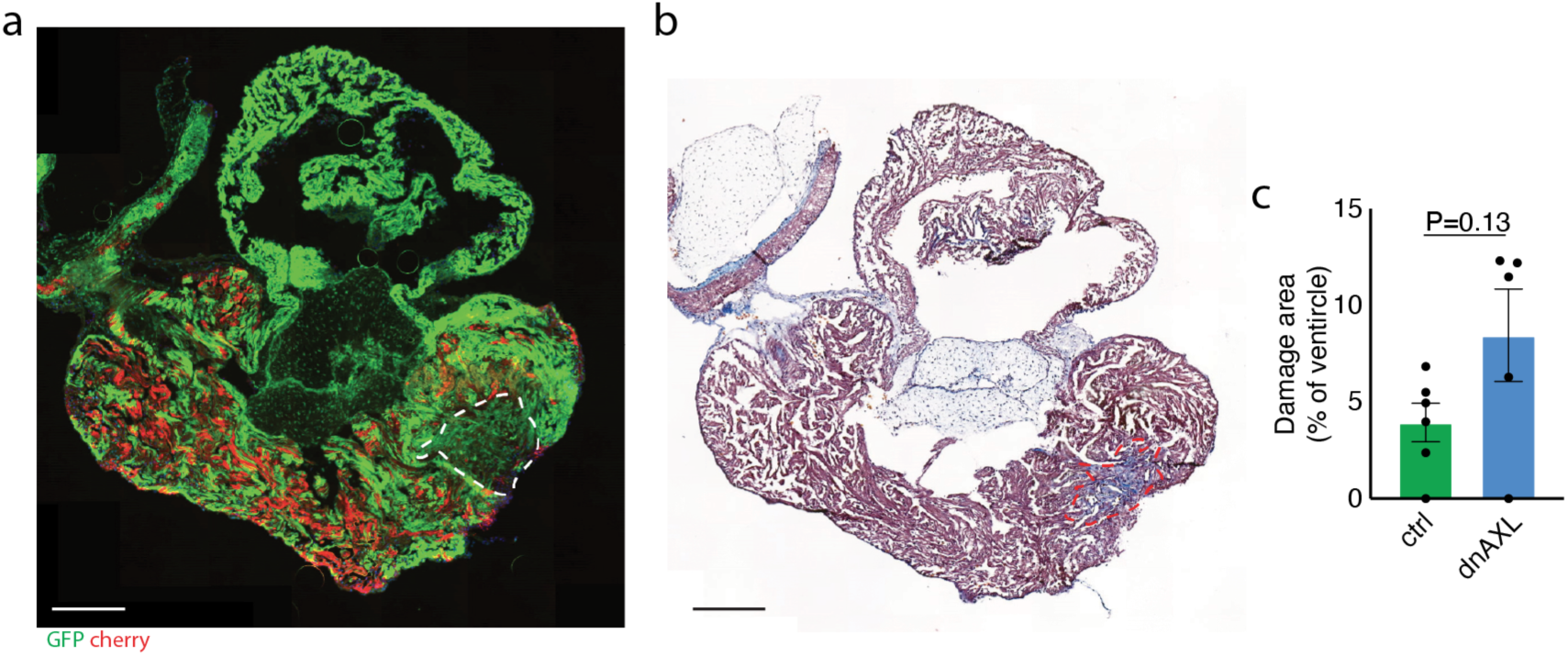
Quantification of injury size in dnAXL animals by fluorescence. **a-c,** Quantification of injury size on similar hearts as seen in Fig. 3 j-m. **a**,**b**, sister sections of ctrl heart axolotl 8 weeks after injury showing transgenic fluorescence (**a**) or Mason’s trichrome staining (**b**). **c**, quantification of injury based on reduction in overall fluorescence intensity of both red and green channels as seen in **a**.

**Figure S11.**
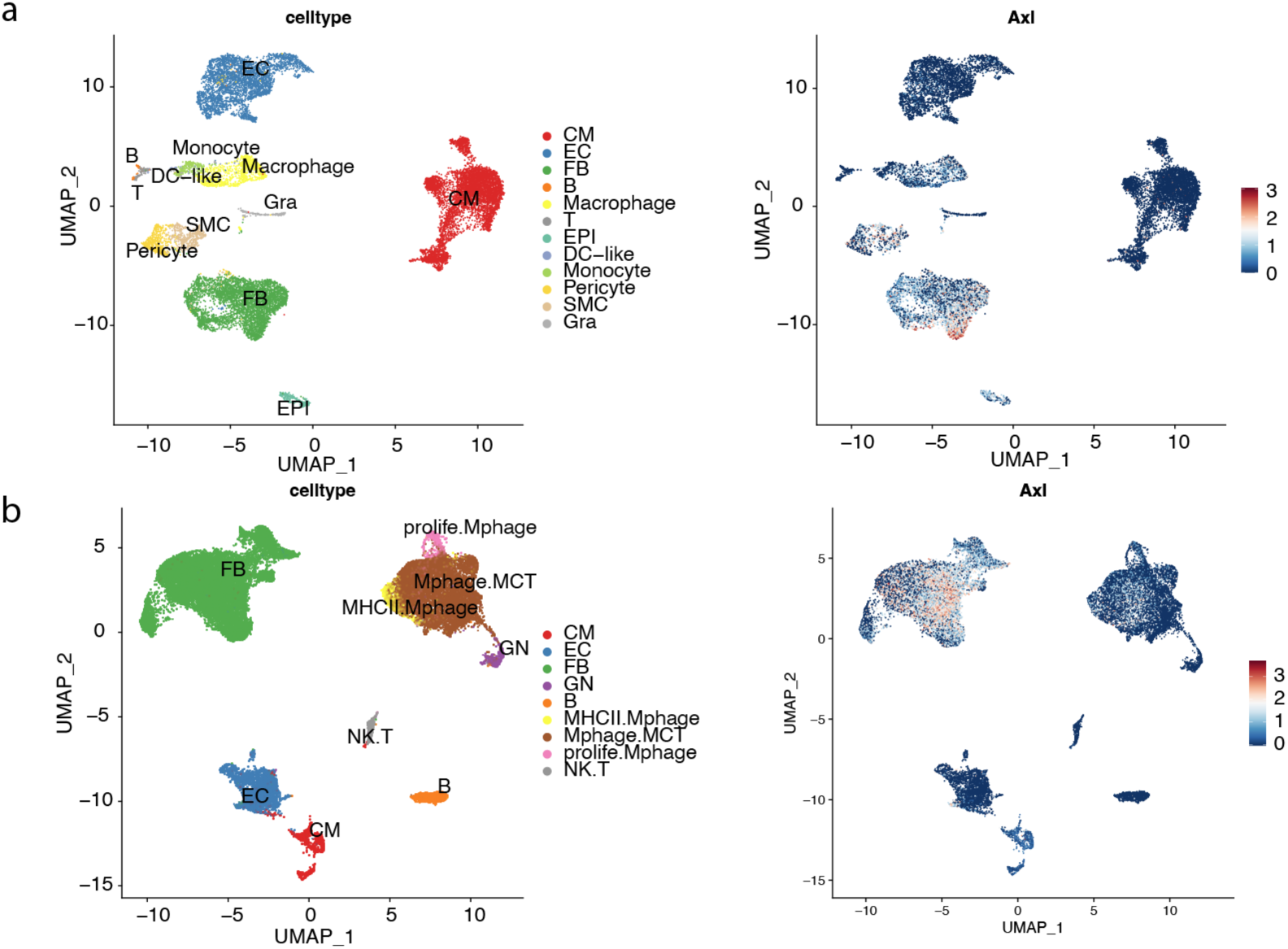
Mouse AXL expression from previously published datasets. **a**,**b** UMAP showing annotated cell clusters re-analyzed from Cui, M. *et al* (*53*) (**a**, left panel) and Hu, P. *et al.* (*65*) (**b**, left panel). Feature plots showing respective AXL expression levels are depicted on the right based on the UMAPs.

**Figure S12.**
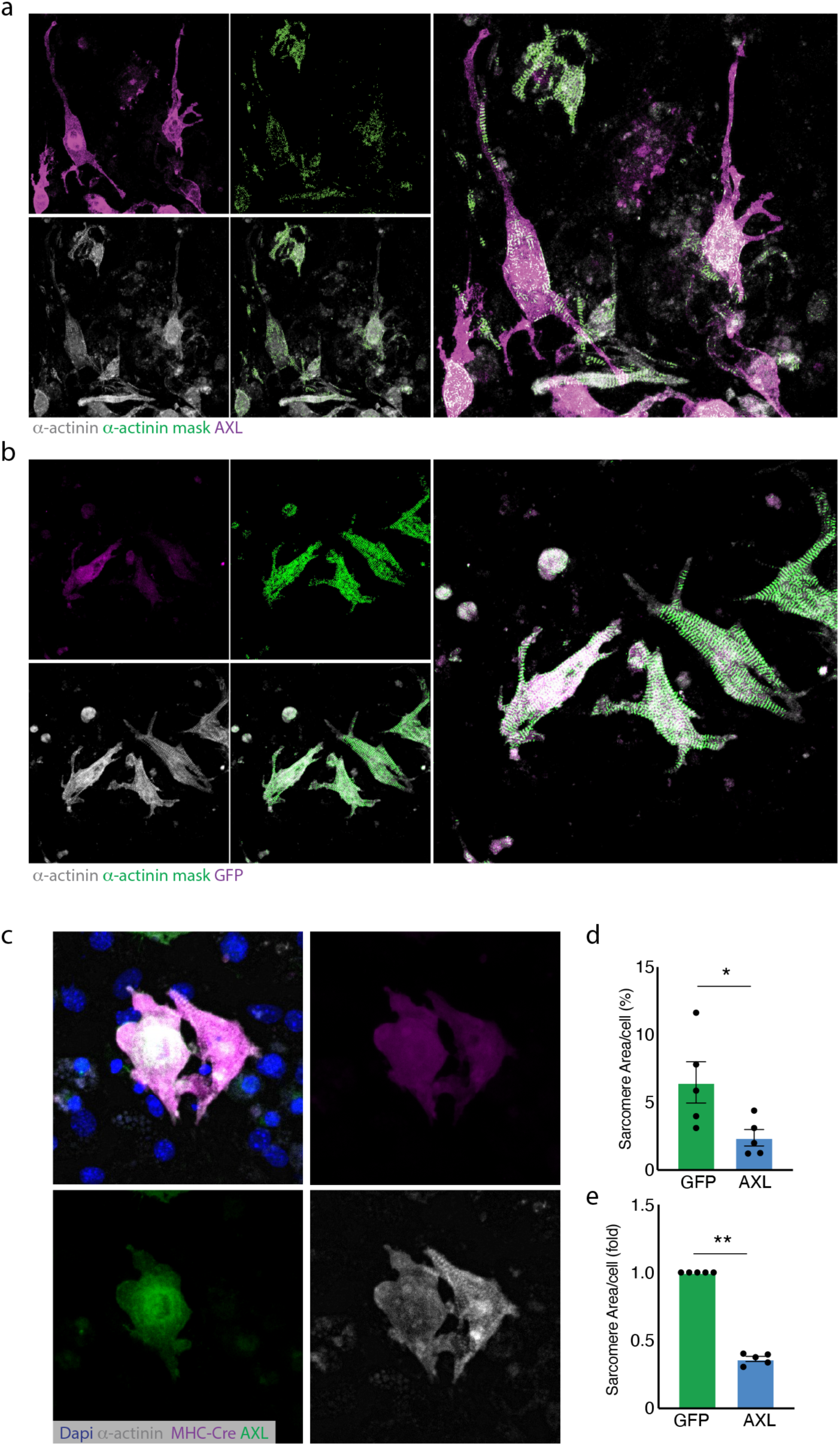
AXL induces sarcomere disassembly in P8 cardiomyocytes. **a,b,** Representative images of AAV9-AXL (a) or AAV9-GFP (b) treated cardiac cultures depicting sarcomere segmentation overlayed on a-actinin and AXL or GFP staining.**c,** representative image of non-AXL OE cell next to AXL OE cell (green) showing disassembled sarcomere structure. **d,e,** Quantification of sarcomere coverage in AXL/GFP OE cells. Sarcomere detection was done in an automated manner and assayed on positive cells detected by Cellpose. Data shown as area/cell (**d**) or fold change relative to GFP ctrl as calculated from a (**e**). Data are presented as average ± s.e.m. \**P* < 0.05, \*\***P** < 0.01 (statistical test: Mann-Whitney two-tailed T-test).

**Figure S13.**
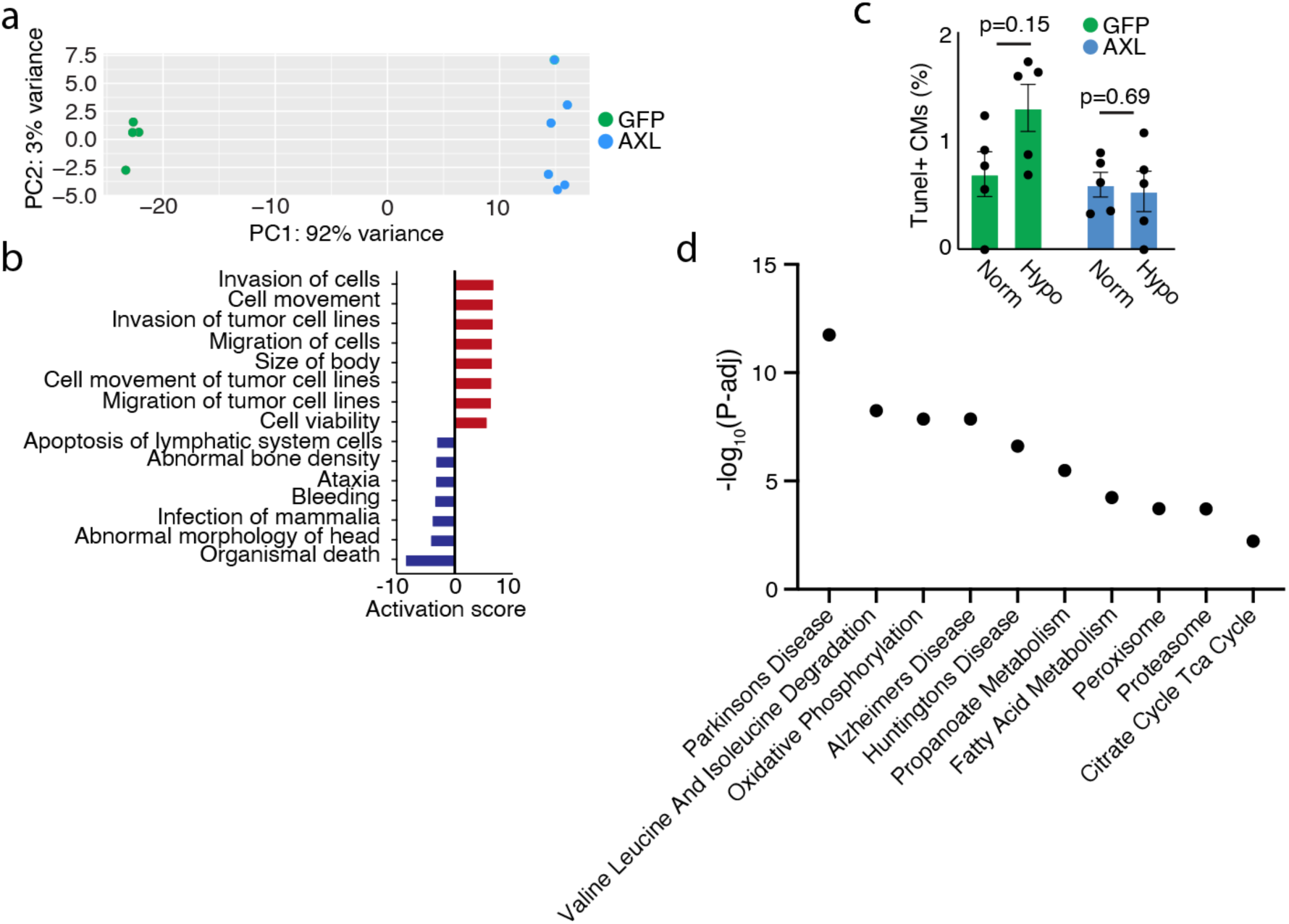
AXL reduces cell death after hypoxia. **a**, Principal component analysis (PCA) of AAV9-AXL or AAV9-GFP infected, bulk RNA-seq samples. **b**, IPA analysis of most activated and inhibited GO terms. **c**, Quantification of cell death of CM by Tunel assay after norm-oxic and hypoxic conditions. statistical test: Mann-Whitney two-tailed T-test**. d**, KEGG enriched go terms based on the bulk RNA-seq results depicted in (**a**)

**Figure S14.**
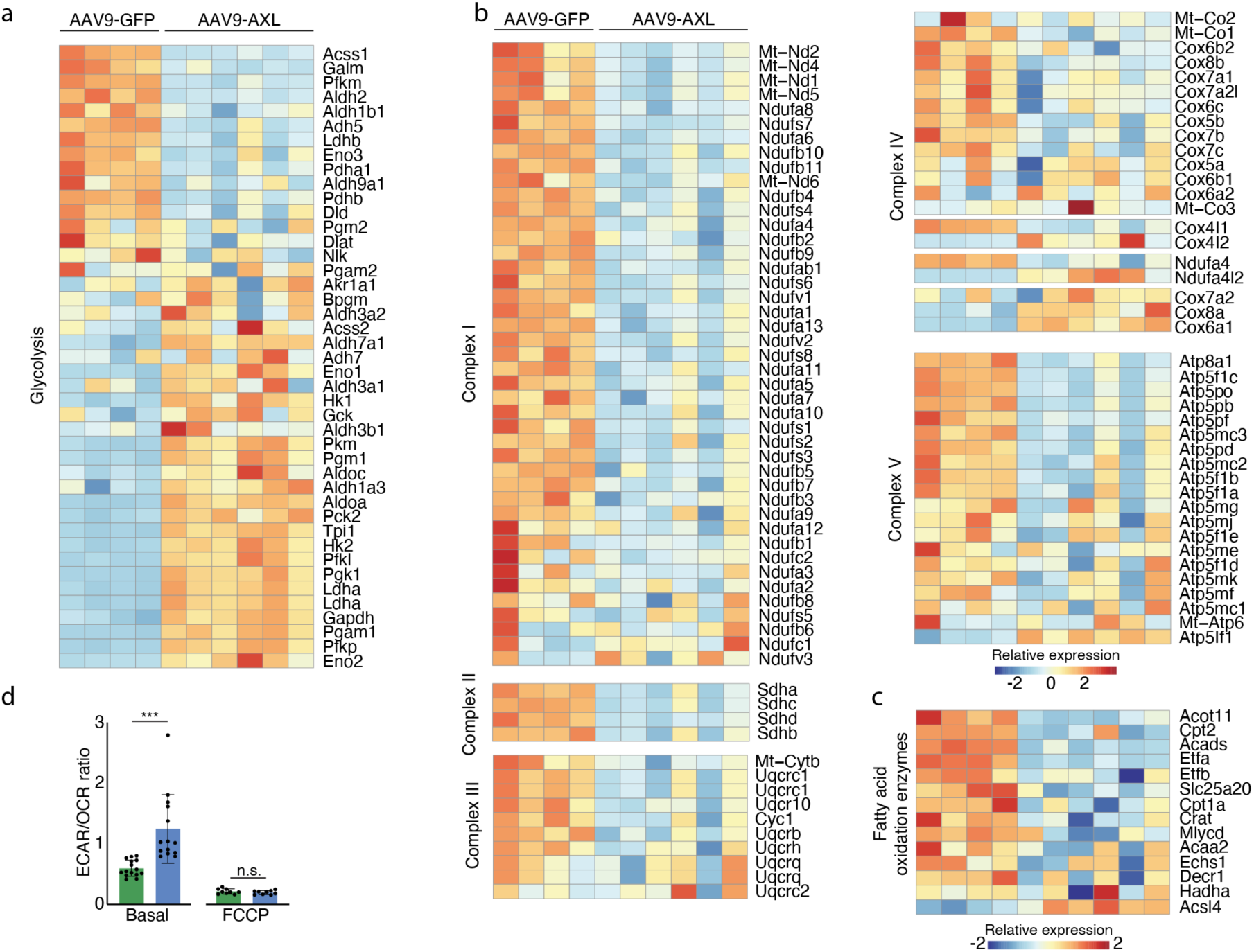
AXL overexpressing CMs upregulate glycolysis related terms and downregulate mitochondrial and fatty acid oxidation genes. **a**, Heatmap depicting glycolysis related gene expression levels. **b**, Heatmap showing all mitochondrial genes from the 5 energy transfer complexes. **c**, Heatmap showing fatty acid oxidation enzymes in GFP or AXL overexpression. Red is increased while blue means downregulated. **d,** Ratio of Extracellular acidification rate (ECAR)/ Oxygen consumption rate (OCR) in the Cell Mito Stress kit, indicating the preference of pyruvate fate of either fermentation or oxidation by mitochondria. Values show either basal levels or with inhibition of mitochondria using FCCP. \*\*\***P** < 0.001 (statistical test: Mann-Whitney two-tailed T-test for basal and FCCP treatment individually).

**Figure S15.**
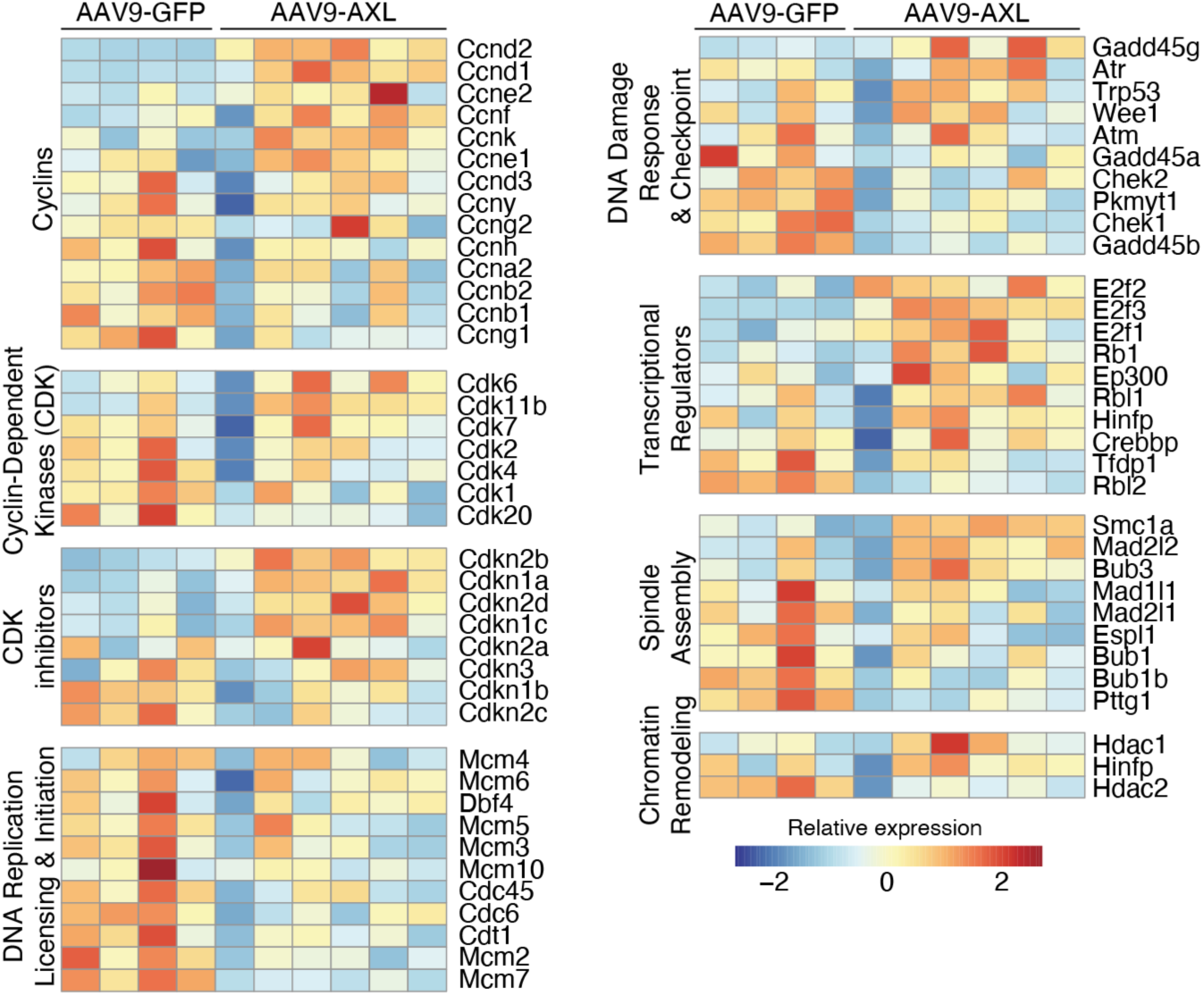
AXL overexpressing CMs show no upregulation of mitosis related genes. Heatmap depicting mitosis related groups and gene expression levels compared to AAV9-GFP ctrl.

**Table S1 | Sequences used for HCR probe generation**

**Video 1-3 | uninjured gCaMP6+ transgenic heart**

Video depicting gCaMP6 fluorescent reporter of the same heart before injury (1), 14 days post injury (2) and 28 days after injury (3). Videos were pseudo colored in Fiji based on intensity using the fire LUT.

## Methods

### Ethics oversight

All axolotl animal experiments were approved by the Magistrate of Vienna (Genetically Modified Organism Office and MA58, City of Vienna, Austria) and the Austrian Ministry of Education, Sciences and Research, under licenses: GZ: MA 58-1432587-2022-12, GZ: 2024-0.438.718, GZ: MA 58-1516101-2023-21, GZ: 2024-0.438.721 and GZ: MA 58-1054623-2020-22, GZ 2024-0.477.503.

### Axolotl husbandry

Axolotls (*A. mexicanum*) were raised in individual tanks in Vienna tap water according to conditions adapted from Khattak et al (*66*). Axolotl matings were performed by the animal-care team at the IMP/IMBA. Axolotl surgery, live imaging and tissue collection were performed under anesthesia in 0.015% benzocaine (Merck, E-1501) diluted with Vienna tap water, using the benzocaine preparation described previously (*66*). Experiments were not blinded, except to the sex of the animals.

### Mouse husbandry

All experiments were performed using mice (Mus musculus) of outbred background from the animal facility of Centro Nacional de Investigaciones Cardiovasculares. These mice were maintained and handled according to the recommendations of the CNIC Ethics Committee, the Spanish laws and the EU Directive 2010/63/EU for the use of animals in research. All procedures were approved by the Ethics Committees of the Centro Nacional de Investigaciones Cardiovasculares and the Regional Government of Madrid (PROEX CNIC-23/20).

### Axolotl transgenesis

The pipeline for axolotl transgenesis were done as previously described by Khattak et al (*66*). Plasmids for axolotl transgenesis were assembled by Gibson Assembly, amplified using Plasmid Maxi Kits (Qiagen, 12163) and verified by whole plasmid sequencing (Plasmidsaurus) before egg injection. One-cell-stage axolotl eggs were sterilized twice for 5 min with about 0.004% sodium hypochlorite solution (Honeywell, 71696) diluted with tap water, then washed well with fresh tap water. Eggs were de-jellied using sharp forceps in 20% Ficoll (Merck, GE17-300-05)/1X MMR/Pen-Strep (Merck, P0781) solution, then held in 10% Ficoll/1X MMR/Pen-Strep solution until microinjection. For microinjections, borosilicate glass capillary needles with filament (Harvard Apparatus, GC100F-15) were pulled using a Flaming/Brown Micropipette Puller P-97 (Sutter Instrument) with settings P = 500, heat = 530 (Ramp test + 30), pull = 100, velocity = 120, time = 150. Then 5 nl of the appropriate injection mix was injected into each de-jellied egg, delivered in two 2.5-nl shots. Egg injections were performed using an Olympus SZX10 microscope using a PV830 pneumatic Picopump (World Precision Instruments) with settings vacuum eject, regulator 25, range 100 ms, timed, duration 10-0. Injected eggs were transferred to 5% Ficoll/0.1× MMR/Pen-Strep solution overnight. The next morning, healthy eggs were transferred to individual wells of a 24-well multiwell plate (Thermo Fisher Scientific, 142475) filled with 0.1× MMR/Pen-Strep solution.

The axolotl lines used in this paper were generated by random insertion I-SceI meganuclease-mediated transgenesis and were screened ±3 weeks post injection, for fluorescent transgene expression using an AXIOzoom V16 widefield microscope (Zeiss).

These lines include:

CAGGs-FUCCI (tgSceI(CAGGs:CDT1[aa1-128]-mVenus-T2A-mCherry-GMNN[aa1-93])^ETNKA^)(*21*), dnAXL-3Xflag-T2a-mCherry (tgSceI(Caggs:loxP-eGFP-stop-loxP-dnAXL-3xFlag-T2a-mCherry)^ETNKA^), Lp-Cherry (tgSceI(Caggs:loxP-eGFP-stop-loxP-mCherry)^ETNKA^), xCMLC2::dER-Cre (tgSceI(xen.CMLC2:TFPnls-T2a-Cre-ERT2)^ETNKA^), IsceI-Caggs::H2b-GCaMP6s. xCMLC2::dER-Cre were crossed either with dnAXL-3xflag-T2a-mCherry for the generation of dnAXL experimental animals or with Lp-Cherry for control. The resulting progeny were screened for whole body flourscent GFP signal (indicative of the dnAXL-3xflag-T2a-mCherry or Lp-Cherry). Positive animals were genotyped for the presence of Cre and those positive were subjected at ±3 weeks post hatching to tamoxifen conversion and induction of dnAXL in the heart. For that animals were bathed overnight in the dark in 4 μM 4-OHT for 3 non-consecutive nights.

### Axolotl cardiac cryo-injury

Cardiac cryo injuries were performed on 8-11 cm animals similar to Godwin J, et al.(*18*) with changes. In short, Animals were placed in 0.015% benzocaine (Merck, E-1501) until pinching of tail or limb shows non-responsiveness. Following this, animals were placed on moist Kimwipes in a supine position and using small iridectomy scissors and forceps, a lateral thoracotomy was performed exposing the chest superficial muscles and cartilage plates. Using two retractors, both cartilage plates and muscle were spread to the sides exposing the pericardial sac. Using iridectomy scissors, the pericardial sac was opened to reveal the beating heart. To perform the cryo lesion, a liquid nitrogen chilled cryo probe was used while the animal thoracotomy was taking place. The tip of the cryo probe was placed on the tip of the ventricle for 5-10 seconds and then immediately washed with saline supplemented with 1x penicillin/streptomycin to release the probe, this process results in a white area indicative of cryo damage. Using forceps, the two retractors were removed and the cartilage plate and muscles returned to original. The skin was closed using surgical glue (Histoacryl, B. Braun) placed directly on the incision and absorbed with a clean Kimwipe. Animals were placed for a 10 min recovery period under benzocaine-soaked towels before returning to clean water containing 0.015% butorphanol with 20% Holtfreters solution with penicillin and streptomycin added to water for 3 days followed by additional 4 days post-op care.

### In-vivo Ca+ imaging and analysis

For Ca+ imaging, axolotls were prepped in a similar way to the cryo injury ending in open chest cavity, exposed heart in a supine position. The entire surgery was performed on a tray which made positioning the animal easier for imaging. Using AXIOzoom V16 widefield microscope (Zeiss) with camera streaming feature enabled, bursts of images were taken to generate the movies of the gCaMP6 fluorescent reporter while the heart beats. Animals were imaged at baseline, experimental midpoint and at endpoint before tissue harvesting. Analysis of injury size and “pseudo-ejection fraction” were done by tracing the outline of the ventricle during systole and diastole. The images were pseudo colored in Fiji based on intensity using the fire LUT. Damaged area showed consistently high intensity area even during diastole, and was measured by outline tracing the intense area. For pseudo-ejection fraction the normal EF equation was used 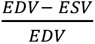, (EDV, end diastolic volume, ESV, end systolic volume) where both volume values were derived from outline tracing of the ventricle.

### EdU administration and detection

Anesthetized axolotls were injected intraperitoneally with 400 μM EdU (diluted in PBS) at a dosage of 20 μl/g. FastGreen dye (Sigma-Aldrich) was added to the injection mix to aid visualization. Injected axolotls were kept out of water for a 10 min recovery period under benzocaine-soaked towels. After recovery, injected axolotls were returned to water.

### AAV production

AAV vectors were designed and constructed using VectorBuilder design studio. Adeno-associated virus serotype 9 (AAV9) particles were produced at CNIC’s Viral Vectors Unit via a dual transfection process in a HEK293T-derived cell line (AAVPro®293T; Takara Bio). This involved the co-transfection of a plasmid containing adenovirus helper sequences, rep (AAV2) and cap (AAV9) genes (pDG9, Plasmidfactory), and a shuttle AAV plasmid.

Following cell lysis in 50 mM Tris (pH 8), 150 mM NaCl, and three freeze/thaw cycles, cells were treated with 150 U/mL Benzonase (Millipore) at 37°C for 30 minutes. Clarification was achieved by centrifugation at 3000 xg for 5 minutes at room temperature.

Viral particles were then purified using iodixanol gradient ultracentrifugation in Polypropylene Optiseal tubes (361625, Beckman-Coulter) at 183 000 xg for 3 hours at 16 °C with a type 70 Ti fixed-angle rotor. Further concentration was performed using Amicon Ultra-15 tubes (100K MWCO, Millipore). Samples were stored in PBS buffer supplemented with 0.001% pluronic F-68. Particle abundance was quantified by qPCR targeting the CMV promoter (primers: ACCATGGTGATGCGGTTTTG and ATGGGGTGGAGACTTGGAAATC)

### Mouse cardiomyocyte isolation

Primary cardiac cells were obtained from P1 or P8 CD1 mouse heart ventricles using Miltenyi’s Neonatal Heart Dissociation kit (130-098-373) in GentleMACS Octodissociator with Heaters according to the manufacturer’s instructions. For cardiomyocyte enrichment, the dissociates were pre-plated in non-coated multiwell plates for 1 hour or further purified using Neonatal Cardiomyocyte Isolation Kit (130-100-825, Miltenyi). Cells were seeded in ibiTreat slides or multiwell plates (81816 or 82426, ibidi) with 1% gelatin coating (G1890, Sigma-Aldrich). Cardiomyocytes were cultured in DMEM (1X) + GlutaMAX without sodium pyruvate (10564-011, Gibco), supplemented with 1% penicillin/streptomycin (P4333, Sigma), 5% inactive fetal bovine serum (F7524, Sigma-Aldrich) at 37 °C and 5% CO2.

In all experiments, AAV9-AXL and GFP were added at a multiplicity of infection of 5E5 VP/cell the day after seeding and left for at least one day before media exchange. In experiments involving administration of AXL inhibitor (R428; HY-15150, MedChemExpress), the cells were treated with the inhibitor or DMSO 24 hours after AAV administration and fixed by day 5 post infection with one media refreshment. All cultures were fixed with 4% paraformaldehyde in PBS for 10 minutes at room temperature before downstream staining procedures. In hypoxia experiments, cells were placed in a hypoxia chamber with 3% O2 for 7 hours the 7^th^ day after infection, then fixed as described inside the chamber to avoid oxygen shock.

### ECAR and OCR measurements

Oxygen consumption rate (OCR) and extracellular acidification rate (ECAR) were measured using a Seahorse XF Pro extracellular flux analyzer with the Cell Mito Stress Test and Glycolysis Stress Test kits (Agilent Technologies). Cardiomyocytes (30,000 cells per well) were plated onto Matrigel Matrix–coated XF Pro Cell Culture Microplates (Corning, 354234) and treated as described. Assays were performed following manufacturer instructions, and data were analyzed using Seahorse Wave software and normalized to cell number using Hoechst 33342 (1:2000) fluorescence. OCR measurements reported mitochondrial oxygen consumption under basal and drug-perturbed conditions, whereas ECAR reflected extracellular acidification arising from both lactate extrusion during glycolysis and CO₂ hydration following mitochondrial pyruvate oxidation. Maximal Respiration was calculated as the maximum OCR after FCCP injection minus non-mitochondrial respiration (after rotenone and antimycin A injection). To assess the metabolic routing of pyruvate between mitochondrial oxidation and fermentation, the ECAR/OCR ratio was calculated for each condition, approximating (Δlactate + ΔCO_’_)/OCR. Increases in the ECAR/OCR ratio were interpreted as a shift toward lactate-producing glycolysis, whereas decreases indicated preferential mitochondrial oxidation of pyruvate. ECAR/OCR values were quantified at baseline and following sequential compound injections in both the Glycolysis Stress Test and the Cell Mito Stress Test to compare metabolic behavior between control and AXL-expressing cardiomyocytes.

### Tissue harvesting

For axolotls, at designated endpoints, animals were placed in 0.015% benzocaine until non-responsiveness to pinching and then handled in a similar way to when cryo injury was performed until the heart was exposed. Using blunt ended forceps, the heart was grabbed at the base of the ventricle and was gently pulled to lift the ventricle and expose the conus arteriosus, then using iridectomy scissors incisions were made to all blood vessels connecting to the heart and then he was removed and washed in 0.7x PBS. For spatial transcriptomics, hearts were moved directly to optimal cutting temperature (OCT) solution and flash frozen in bath full of Isopentane chilled with liquid nitrogen following by moving to -70^0^C till sectioning. For histology, hearts were moved to overnight fixation in 4% formaldehyde solution following by 5 days in 30% sucrose prior to embedding in OCT and freezing in Isopentane.

For mouse, animals were sacrificed in a CO2 chamber. After securing the body, the chest was open with surgical scissors to expose the liver and the chest cavity. Three cuts in the liver were made to allow blood drainage. The left ventricle wall was exposed using forceps and then perforated with a 23G needle placed in a 10 mL syringe. Approximately 10 mL of PBS with heparin were slowly introduced, followed by 10 mL of KCl 50mM solution with heparin to stop the heartbeat. Finally, 10 mL of PFA 4% of histological quality were passed through. After perfusion, the heart was carefully removed, cleaned and placed in PFA 4% overnight, tilting at 4°C. 24 hours after, the PFA solution was removed and 30% sucrose in distilled water was added for at least 24 extra hours before mounting in OCT.

### HCR probe design

HCR probe pairs were designed against unique mRNA sequences of the candidate gene. BLAST alignment of the gene sequence against the axolotl transcriptome Amex.T_v47 was used to identify unique sequences. Unique 52-bp sequences were then inputted to R studio software and using the Statistical Patterns in Genomic Sequences (SPGS) library the unique sequences were cut in half and an HCR amplifier sequence was added. These edited sequences were ordered as Oligo opool (IDT, 50pmol). All buffers including amplification buffer, as well as HCR hairpins were purchased from Molecular Instruments. Detailed sequences of each probe can be found in table S1.

### Tissue sectioning and staining

Tissue sectioning was performed as serial sections spanning 10 slides with 1 skip between slides, hence each slide contained sections from the entire heart. Sectioning were done using Leica CM3050 S Research Cryostat and thickness was 10µm in all immunofluorescent or immunohistochemistry experiments.

Masson’s Trichrome staining was performed using the Masson Trichrome kit (cat. 04-010802, Bio-Optica) with the following modifications: The mix of reagents A and B was left to act for 6 minutes (instead of 10). Reagent D was left to act for 2 minutes (instead of 4). Reagent F was left to act for 3 minutes (instead of 5). Dehydration was done for 10sec in 95% alcohol and 1 minute in absolute ethanol. Slides were air-dried and mounted with xylene-based mounting medium.

For immunofluorescence of axolotl tissues (apart from EdU staining), slides were allowed to equilibrate at room temperature in a closed slide box for 30 minutes and then the region of interest was surrounded with hydrophobic pen. Slides were washed with PBS to remove OCT and then were permeabilized at room temperature with 0.5% triton x100 diluted in PBS for 15 minutes. Next, slides were blocked with 3% BSA and 0.1% triton x100 diluted in PBS for 30 minutes. Slides were placed in a humid chamber on which primary antibody (MF20, DSHB, 1:200 - 3X DYKDDDDK (flag) Tag, Cell Signaling technology, 1:200) diluted in the blocking solution, was added to the slide and covered with parafilm and placed in 4^0^C. The following day, the slides were allowed to equilibrate for 1 hour at room temperature and then were washed with PBS, 3 times. Following this, the slides returned to the humid chamber and incubated with secondary antibody for 2 hours. DAPI was included in the secondary staining solution at a concentration of 10 μg/ml. Slides were washed well with PBS and mounted in Abberior Mount liquid antifade mounting media (Abberior) for imaging.

For immunofluorescence of mouse tissues Axl (PA5-106118, Thermo Fischer) and α-Actinin (A7811, Millipore) staining slides were allowed to equilibrate at room temperature in a closed slide box for 30 minutes. Slides were washed 3x with PBS, 3 minutes each to remove OCT and then placed in a humid chamber. Slides were incubated with 2% BSA 0.1% Triton in PBS for blocking and permeabilization for 1 hour. Then they were incubated with primary antibodies (Axl 1:200, Actinin 1:200) diluted in the same blocking solution at 4^0^ overnight. The next day the slides were washed with PBS three times for three minutes. Following this, the slides returned to the humid chamber and incubated with secondary antibody for 2 hours. DAPI was included in the secondary staining solution at a concentration of 10 μg/ml. Slides were washed well with PBS and mounted with DAKO Antifade mounting medium.

EdU staining was done according to the manufacture’s protocol (Thermo Fisher, C10340) and then followed by the normal immunofluorescent staining protocol.

For HCR, staining was done according to manufactures protocol for fixed frozen tissue sections (Molecular Instruments), without post-fixation or Prot-K steps. All buffers and staining reagents used were bought from Molecular Instruments. Slides were counterstained with Dapi and in some cases with Phalloidin-Alto 594 (Sigma, 51927). Slides were mounted in Abberior Mount liquid antifade mounting media (Abberior) for imaging.

Mason’s trichrome images were taken with the Pannoramic 250 FLASH III Digital Scanner (3DHistech). Flourscent images were taken either by Pannoramic 250 FLASH III Digital Scanner (3DHistech) for EdU or with Olympus IX3 Series (IX83) inverted Spinning Disk Confocal microscope with CellSense software (Olympus). HCR images were acquired with an LSM980 AxioObserver inverted confocal microscope with ZEN software (Zeiss) with AiryScan 2.

### Imaging quantification

After acquisition EdU labeled images were exported to Fiji (Imagej) and then were counted in the 300um area adjacent to the injury zone. In the WT animals, CM were detected by staining for MF20. In the ctrl group to the dnAXL, CM were selected based on the presence of mCherry and loss of GFP fluorescence, only cells which lost GFP completely were used. In the dnAXL CMs were selected based on the staining for Flag-tag and complete loss of GFP signal. HCR images were processed using the ZEN blue software (Zeiss) batch AiryScan processing tool. For quantification of cell numbers (i.e. number of positive/negative cells), events tool in ZEN blue was used and cells were manually counted for the presence/absence of signal. Quantification of *Nppa* and *Ankrd1* in dnAXL animals was performed by counting positive/negative HCR signal in cells without GFP (converted) adjacent to the injury zone (up to 3 rows of cells). The cells selected showed an elongated morphology with the presence of a dapi positive nuclei. The Lp-cherry ctrl animals were chosen by the presence of the mCherry signal however mCherry signal in dnAXL was too low to detect even when performing antibody staining thus cells were chosen for lack of GFP and elongated morphology alone. Mean gray values, region of interest was exported from ZEN to Fiji (ImageJ). For mean gray value in a specific line (as done agbl1 quantification), thick line plots were generated for 300um going from the epicardium into the middle of the ventricle and the values from the plot were exported. Next, the values were “binned” to 10-100,100-200,200-300um by averaging all values in the range. For measuring individual HCR signal spots, the ROI was masked by setting a threshold that allowed good signal to noise and counted using analyze particle’s function. The ROI area was measured to normalize particles/area.

Analysis of injury size was performed on images acquired from the slide scanner of tissue sections stained with Mason’s trichrome or using fluorescence derived from the inserted transgene. All hearts were positioned in the same way, producing similar section orientation. For the stained sections the 3 sections (out of 10-12 per animal) who showed the largest damage area by eye were selected for analysis. Damage area was measured using Case Viewer software (3DHistech), and identified as general abnormal morphology such as: void area, unmuscularized area, area positive for fibrotic (blue) staining or pale staining color, as was highlighted by Dittrich A, et al(*19*). For each section, measurement of the entire ventricle area was also performed for calculation of damage % from total ventricle area. Adjacent sections from the same hearts were also imaged for presence of fluorescence signal and damaged areas counted by trichorme staining were validated to have lower intensity of fluorescence.

### Cell culture immunostaining and imaging

After fixation, cultured cells were permeabilized with 0.5% Triton X-100 in PBS for 10 min, and blocked with universal TNB blocking reagent (FP1012, Perkin Elmer) for 1 h at room temperature. For immunostaining, the cells were incubated overnight at 4°C with the following antibodies diluted in the blocking solution: AXL (1:100, PA5-106118, ThermoFisher), Alpha actinin (1:300, ab9465, abcam), GFP (1:200, R1091P, Origene).Cells were then washed three times with PBS and stained for 60 min at room temperature with suitable secondary antibody, diluted 1:500 in PBS-Triton 0,1%, and DAPI 1:1000 (4,6-diamidino-2-phenylindole dihydrochloride). If cells needed to be stained for tdTomato for linage tracing, they were then washed three times with PBS, blocked with 5% Normal Rabbit Serum (011-000-120, Jackson ImmunoResearch) in PBS-Triton 0,1% for 30 minutes at room temperature and stained with anti RFP-CF594 conjugated antibody (1:200, 20422, Biotium) for 1 hour at room temperature. Finally, cells were washed three times with PBS and kept in PBS-Azide 0,1% until imaging.

For assessing cell death in the hypoxia experiment, TUNEL staining was performed using Terminal Transferase kit (3333566001, Roche) and Biotin-16-dUTPs (11093070910, Roche). After permeabilization, cells were incubated with buffer TdT 1X CoCl2 1mM for 30 minutes at room temperature. Then, TdT reaction mix (buffer TdT 1X CoCl2 1mM + 0,2% biotin-16-dUTPs + 0,3% Terminal transferase) was incubated for 1h at 37°C in humid chamber. Immediately, the reaction mix was replaced with citrate buffer 10mM (pH 6) in PBS to stop the reaction for 15 minutes at room temperature. After three washes with PBS, the protocol for immunofluorescence described above was followed, including Streptavidin 647 (1:500, S21374, Life Technologies) in the secondary antibody mix.

For cell cycle entry assays, Click-iT™ EdU Alexa Fluor™ 647 Imaging Kit (C10340, ThermoFisher) was used according to manufacturer’s instructions. After EdU detection, the protocol for immunofluorescence described above was followed.

Microscopy was performed using a Leica SP8 Navigator at the Microscopy & Dynamic Imaging Unit, CNIC, co-funded by MCIN/AEI/10.13039/501100011033 and FEDER“Una manera de hacer Europa” (#ICTS-2018-04-CNIC-16).

### Automated sarcomere segmentation

For *in-vitro* quantification, all image analyses were performed in Fiji/ImageJ using custom macros developed in-house, incorporating the Bio-Formats importer, Cellpose 3 plugin, and standard ImageJ functions. Multichannel confocal stacks were imported and analyzed in batch mode to ensure consistent processing across all samples.

To determine the most in-focus z-slice for each image, the mean intensity across all slices was calculated, and the slice with the highest mean signal was selected for downstream measurements. Images were rescaled twofold in the x–y dimension to standardize pixel sampling across datasets.

Sarcomeric structures were segmented using a custom orientation-independent fast Fourier transform (FFT)-based filter that enhances periodic banding patterns regardless of orientation. For each slice, an eight-angle (22.5° steps) rotated frequency-space mask was applied to the sarcomere channel, followed by forward and inverse FFT transformations to isolate oriented signal components. The resulting images were concatenated, thresholded (adjusted per experiment), and filtered (median radius = 2) to generate a binary mask representing the total sarcomeric area.

Cells were segmented from the cytoplasmic channel using Cellpose 3 (GPU-accelerated, custom-trained model). Prior to segmentation, the channel was smoothed. Label images generated by Cellpose were converted to individual regions of interest (ROIs) within Fiji, followed by filtering to exclude objects smaller than 500 µm² or with a low mean intensity. The resulting ROI set was overlaid on the source image for quality control.

Quantification of sarcomeric signal per cell was performed by combining Cellpose-derived ROIs with the FFT-based sarcomere masks. For each ROI, the most in-focus slice was identified as described above, and the mean pixel intensity within that region was measured from both the raw sarcomere mask and a smoothed version. Thresholding (adjusted per experiment) and morphological closing (five iterations, three counts) were applied to generate conservative binary representations of sarcomeric area. For each cell, the mean pixel value within the ROI on the sarcomere mask was defined as *Percent of sarcomere Area*, and the mean value on the smoothed mask as *Percent Merged Area*.

*In-vivo* quantification was performed as described above with the exception that regions of interest (ROIs) were selected manually based on AXL+/- and α-actinin+/- regions. Selected ROIs were taken from elongated cellular morphology that showed aligned sarcomere staining. ROI selection was performed to select roughly similar sizes for each group.

### Spatial transcriptomics

Molds containing frozen hearts from 0 (non-injured),1, 4, 7 and 14 post-injury (as seen in Tissue harvesting section) were placed in cryostat (Epredia CryoStar NX70 cryostat, Thermo Fisher) until equilibration and were sectioned at 10μm thickness. Blocks were cut and sections were placed on SuperFrost plus slides (Thermo Scientific, 630-0951) and were rapidly stained with H&E (quick wash with H2O, 15sec in Hematoxylin (Fisher Scientific, 6765002), quick wash with H2O, counterstain for 15sec with Eosin (Fisher Scientific, 10483750) and wash with H2O before mounting the slide) and evaluated for localization and extent of damage. This process was repeated until the optimal region was reached.

To assess optimal permeabilization conditions for Visium experiment, 8 adjacent sections were placed on the Visium Spatial Tissue Optimization Slide (10X Genomics, 1000193) and were subjected to different permeabilization times. Optimal permeabilization time was 30mins and was used in the Visium experiment.

Once all sections were selected for Visium, blocks were placed in cryostat until equilibration along with the Visium spatial gene expression slide (10X Genomics, 1000187). Blocks were then aligned and a single 10μm section was placed on the Visium slide. After all sections were placed on the Visium slide, the downstream pipeline was continued in accordance with the manufacturer’s guidelines (10x genomics, Visium Spatial Gene Expression Reagent Kits User Guide). The slides were imaged with Axio Imager.Z2 (upright) with Zeiss Axiocam 506 Color camera. Visium data was processed with SpaceRanger (v1.3.0) with the images manually aligned, with reference genome mexG_v6.0-DD and transcriptome assembly AmexT_v47 (*67*). The filtered feature barcode matrix of each slide was imported to Seurat (v4.1.1) and merged.

### Nuclei isolation

To obtain single nuclei, 6 hearts per time point were wrapped in aluminum foil and were flash-frozen in liquid N2, after which they were moved on dry ice to -70 until use. Working in the +4^0^C room, the tissues were transferred from dry ice into Dounce tissue grinder set (Merck) containing 2ml of Nuclei PURE Lysis Buffer (Sigma Aldrich, Cat# L9286) supplemented with 0.1% triton x100 and 1mM dithiothreitol (DTT), 1× protease inhibitor (cOmplete Mini, Merck) and 0.8 U/μl RNAse inhibitor (RNaseOUT, Thermo Fisher). After 10 strokes using the loose pestle (A) and additional 6 with the dense pestle (B), the homogenate was filtered through 70 μm cell strainer and then 40 μm cell strainer (Corning). After centrifugation (300g, 5 min, 4 °C), the supernatant was removed and the pellet was resuspended in Nuclei PURE Storage Buffer (Sigma Aldrich, Cat# S9183). After additional centrifugation step (300g, 5 min, 4 °C), the majority of supernatant was removed and the nuclei were resuspended in residual volume. Nuclei were stained with 7-AAD viability staining solution (BioLegend) and Dapi, and then assessed using NucleoCounter NC250 for number and integrity.

### Multiomic sequencing

For snRNAseq and ATACseq profiling, 16k nuclei per channel (to get a recovery of ±10k nuclei) of a 10x microfluidic chip device. Combined snRNAseq and snATACseq were generated with the Chromium Single Cell Multiome ATAC + Gene Expression kit following manufacture recommendations (10x Genomics). Final libraries were sequenced on Illumina NovaSeq 6000 S4 lane 300 cycles.

### snRNA-seq processing and analysis

#### snRNA-seq processing and quantification

The UMI counts of each snRNA-seq library from 10x multiome were quantified with the same processing pipeline as in (Lust et al., 2022), Briefly, the raw FASTQ files of snRNA-seq were processed on the latest axolotl transcriptome annotation (v47) based on the genome release (v6.0) with kallisto (v0.46.0) and bustools (v0.40.0) with default parameters. To account for the large number of reads from intronic regions in axolotl genome due to nuclei isolation, both exonic and intronic regions were considered to quantify the UMI counts of genes.

#### Cell filtering and normalization

With the UMI counts of genes in each library, the empty drops were first detected using the DefaultDrops function with expected cell number of 8000 in R package DropletUtils (v1.14.2). The gene expression matrix and sample information were then imported to Seurat (v4.1.3) and further filtered if nFeature_RNA < 200 or nFeature_RNA > 10000 or percent.mt < 60. To discard the detected doublets in the scRNA-seq data, the DoubleFinder (v2.0.3) approach were applied and only singlets were kept. The cleaned UMI counts were normalized using LogNormalize method with scale.factor = 10000 and highly variable genes were identified with the “vst” method. Moreover, the cell-cycle scores and phases were assigned with CellCycleScoring in Seurat using the known G2/M and S phase markers.

#### Annotation of major cell types and subtypes

To annotate the different cardiac cell types, using the Seurat package (v4.1.3) we first ran “low resolution” clustering to obtain small number of populations and detect main cell types. Once satisfactory clustering was achieved, using the FindAllMarkers function, we identified marker genes for each main cluster which allowed us to broadly define the clusters (i.e. CM, EC, FB etc.). Next, each main cluster was sub clustered with different iterations to a point that each sub-population had still unique markers compared to other cell types within the same cluster. Next, we defined marker genes for each sub-population within the context of the main population and we began annotation based on key known markers. Cells which possessed known markers from other main populations, such as a CM with EC canonical marker expression, were labeled as possible doublets and were filtered out. Although this approach would limit the ability to identify “rare” cell populations, as this is a first-of-its-kind atlas and likely the basis for future dataset annotations we opted for higher confidence nuclei annotations. Once all possible doublet and low quality cells were filtered out, sub-clusters were overlayed onto the Visium slide and thus annotated both by known marker genes as well as spatial localization.

#### Trajectory inference with PAGA velocity graph

To calculate RNA velocity for the snRNA-seq data, we first applied the kallisto-bustools pipeline as in (Lust et al., 2022) to quantify both spliced (i.e. from exon) and unspliced (from intron) count matrix. Dynamical model was used for the RNA velocity computation in scvelo (v0.2.5). To infer the trajectory among previously annotated subtypes of CMs and ECs, the PAGA graphs with velocity-guided directionality were calculated and visualized in original UMAP embedding of snRNA-seq data (i.e. quantification with both exons and introns).

### snATAC-seq data processing and analysis

snATAC-seq data and scRNA-seq were simultaneously collected from the same cell in according to 10x multiome protocol. The paired snATAC-seq and snRNA-seq data from the same sample were processed and quantified together with cellranger-arc (v2.0.0), as in (Lust et al., 2022) using the fragmented contigs of axolotl genome (v6.0), to overcome the hurdle due to the large genome size. Here we kept also only the snATAC-seq output from the cellRanger-arc. Moreover, prior the snATAC-seq data analysis, we first combine peaks identified from each sample to have a list of consensus peaks. We then performed the quantification for consensus peaks in Signac (v1.9.0) for cells that are found in snRNA-seq. Cells were further filtered if they have less than 200 or more than 100000 detected features, or with Seurat parameter nucleosome_signal > 6 or TSS.enrichment < 1). The snATAC-seq modality was integrated with the snRNA-seq previously quantified using kallisto-bustools pipeline. The cell type and subtypes annotations of snRNA-seq were transferred to snATAC-seq data; the low-dimensional UMAP embedding of snRNA-seq was also used for visualization. The motif activities were inferred using chromVAR(v1.16.0).

### Spatial transcriptomics (ST) processing and analysis

#### ST processing and quantification

The raw FASTQ data collected from axolotl hearts at different injury stages were processed with the similar kallisto-bustools pipeline as previous snRNA-seq data processing (Lust et al., 2022). In addition, gene expression was quantified using the axolotl exon annotation (v47) and H&E images of Visium slices were manually aligned using 10x Loupe Browser (v8.0.0). The gene expression matrix, spot barcodes and associated images were imported to Seurat (v4.3.0) and data from different time points and biological replicates were merged into one single SeuratObject (v4.1.3). Furthermore, we filter spots with low quality detection if nCount_Spatial < 300 or nFeature_Spatial < 200. Lastly, the normalized expression of all detected genes was obtained using SCTransform method (v0.3.5) with 3000 highly variable genes.

#### Manual segmentation of axolotl heart in H&E image

To facilitate the downstream analysis, we segmented manually the ventricle regions across all samples based on the H&E morphology and marker genes using SPATA2 (v2.0.4); at each slice, we depicted also manually the injury region (IR), border zone (BZ) and remote zone (RZ) at day 1, 4, 7 and 14 and two intact regions at day 0.

#### Cell type decomposition in Visium spatial data

Multiple cell types/states can be located in a single Visium spot. Therefore, we leveraged our snRNA-seq data to infer the cell type decomposition at each spot using RCTD method implemented in spacexr (v2.0.1). To increase the decomposition accuracy, time-specific cell subtypes identified from snRNA-seq were provided for RCTD with double mode and the output were normalized into subtype proportions (i.e. sum is 100%). To visualize the subtype proportions, the SpatialPlot function from Seurat (v4.3.0) was modified and customized.

#### Neighborhood enrichment analysis

To assess and compare the spatial dependencies between cell subtypes in different regions (i.e. intact region, IZ, BZ and RZ), we applied the multi-view model in mistyR (v1.7.0) for each slide by taking the corresponding subtype decomposition from RCTD as input, in which all views (i.e. intrinsic, juxta and para) were considered. To summarize the dependence scores, we first excluded the negative dependencies (mutual exclusion between cell types) and calculated the median values of scores from multi-views in BZs if they are larger than those in intact regions and RZs. To visualize the cell-type pairs enriched in the BZ neighborhood, the netVisual_circle function from CellChat (v2.1.2) was modified and customized.

#### Cell-cell communication analysis

To predict the potential ligand-receptor interaction in BZs of axolotl regeneration, we specified time-dependent source and receiver cell subtypes and performed cell-cell communication analysis with LIANA (v0.1.13) that systematically aggregate the analysis results from multiple data resource (“Consensus”, “CellPhoneDB” and “CellChatDB”) and also four different statistical methods (“natmi”, “connectome”, “logfc” and “sca”).

#### Ligand-receptor interaction visualization

To visualize the predicted ligand-receptor interaction from LIANA, we modified and customized the CircosPlot function from Connectome (v1.0.1). To further visualize individual ligand-receptor interaction on top of image, we imputed the Visium data by levering our snRNA-seq data and inferred cell decomposition and visualized the interaction score using NICHES (v1.0.0).

### Processing of published sc/snRNA-seq datasets in neonatal and adult mouse

The filtered matrix and barcodes of sn/scRNA-seq data in neonatal mouse from GSE130699 (Cui et al., 2020) (*53*) and GSE153480 (Wang et al., 2020) (*68*) were downloaded. These datasets were processed and analyzed together in Seurat using the cell type annotations provided by (Cui et al., 2020) (*53*) and (Miyara et al., 2025) (*69*). We applied the RPCA method in Seurat to integrate those two datasets from two research groups. Similarly, for sn/scRNA-seq data in adult mouse, the filtered matrix and barcodes of CMs from GSE120064 (Ren et al., 2020) (*70*) were downloaded and only the wild type mouse data at week 0 and 2 were selected. For interstitial cell annotation *scRNA-seq from (Forte et al., 2020)* (*71*) was used and annotated using known markers. We performed again the RPCA method to harmonize those two datasets.

### Bulk RNA-seq library prep and pre-processing

Purified neonatal cardiomyocytes were infected one day after seeding. The cells were lysed and scraped 3 days after infection using RNeasy micro kit (74004, Qiagen) and processed according to manufacturer’s instructions.

NGS experiments were performed in the Genomics Unit of CNIC. For library preparation, NEBNext Multiplex Oligos for Illumina (Index Primers Set 2) (E7500, New England Biolabs) was used. Sequencing was performed using Illumina NextSeq 2000 Sequencing System with a P2 flow cell (1 × 100 bp, 100 cycles, ∼400 million reads).

Generated reads were trimmed for adapters and bad quality sequences using Trimmomatic (v0.39) with the parameters ILLUMINACLIP:TruSeq3-SE.fa:2:30:7 SLIDINGWINDOW:4:20 MINLEN:50. Trimmed reads were then aligned to the mouse genome (GRCm39) using HISAT2 (v2.1.0) with default parameters. Lastly, reads were quantified by using the featureCounts subpackage from subread (v2.0.2) against the annotation for the same version of the genome.

### DEG detection in AAV9-AXL bulk RNA-seq data

We detected DEGs between AXL and GFP treated groups by running the Deseq2(*72*) package in RStudio for statistical analysis. Genes with less then 30 reads across all samples were filtered out and subsequently normalized. Following this, the differential expression analysis was performed using DEseq2 (v1.42.1).

We used adjusted p-value <0.05 as significance threshold. PCA analysis of our acquired 6 AAV9-GFP and 6 AAV9-AXL samples showed that all AXL samples clustered together while 2 GFP samples did not cluster with the other 4 and thus they were excluded from analysis.

## Data and materials availability

All code used for raw data processing and statistical analysis is available at https://github.com/labtanaka/heart_regeneration. Sequencing Fastq files have will be available after peer review. All other data is found in the figures of the paper.

